# The paraspeckle protein NONO potentiates the antiviral innate immune response through chromatin regulation

**DOI:** 10.64898/2026.07.11.737985

**Authors:** Adam Hage, Mikhaila Janes, Byron Shue, Tovah E. Markowitz, Suhyeon Yoon, Jeffrey G. Shannon, Paul A. Beare, Rebecca M. Broeckel, Justin B. Lack, Craig Martens, Sonja M. Best

## Abstract

Type-I interferons (IFN-I) and IFN-stimulated genes (ISGs) are central to antiviral defense, while dysregulation can drive autoimmunity. *IFNB1* expression is controlled by a highly ordered multiprotein complex composed of IRF3/7, NFκB, and ATF2/c-Jun (AP-1) that recruit coactivators and chromatin-remodeling proteins to expose the *IFNB1* promoter for the RNA polymerase II (RNA Pol II) transcriptional machinery. Here, we identified the paraspeckle protein non-POU domain-containing octamer-binding protein (NONO) as a critical facilitator of innate immune activation. Loss of NONO enhanced replication of multiple orthoflaviviruses including West Nile virus due to impaired induction of IFN-I and ISGs. NONO did not affect upstream signaling but instead promoted chromatin accessibility and promoter access for RNA Pol II to drive expression of *IFNB1*, ISGs, and proinflammatory cytokines. These findings position NONO as a key regulator of antiviral gene expression and reveal chromatin-levels of control that determine effective antiviral immunity.

## Introduction

Infection by pathogenic viruses triggers the induction of antiviral type-I and III interferons (IFN-I/III), interferon-stimulated genes (ISGs), as well as proinflammatory cytokines and chemokines that coordinate to limit virus replication, recruit immune cells, and establish an antiviral state in tissues. Following infection, viral nucleic acids (RNA and DNA) can be highly immunostimulatory through ligation of pattern recognition receptors (PRRs) to initiate these cell-intrinsic responses. Major cytosolic PRRs that respond to viral infection resulting in expression of antiviral IFNs include the retinoic acid-inducible gene-I (RIG-I)-like receptors (RLRs), cyclic GMP-AMP synthase (cGAS), and interferon-gamma-inducible protein 16 (IFI16). The signaling cascades downstream of different PRRs converge through activation of latent transcription factors (TFs) including IFN Regulator Factor 3 and 7 (IRF3 and IRF7), Nuclear Factor κB (NFκB), and Activator Protein 1 (AP-1). Phosphorylation of IRF3 and IRF7 drive their translocation to the nucleus to initiate production of IFN-α and IFN-β. IFN-α/β are key cytokines that signal through the IFNAR1 and IFNAR2 receptors to activate the Janus kinases (JAK1 and TYK2) responsible for phosphorylating Signal Transducer and Activator of Transcription 1 and 2 (STAT1 and STAT2) that then assemble together with IRF9 into the transcriptionally active ISGF3 complex. Following its translocation to the nucleus, ISGF3 drives further expression of several hundred ISGs. Collectively, this transcriptional response underpins comprehensive defenses against invading organisms. However, dysregulation of IFN signaling can promote immune-mediated pathology following infection or sterile forms of autoimmunity. Although the importance of master TFs like IRFs and STATs for IFN-I and ISG responses has been firmly established, the breadth of gene targets, temporal changes in gene expression, and cell-specific transcriptional programs strongly implicate involvement of additional factors whose contributions are underinvestigated.

Proper regulation of innate immunity requires strict control of gene expression to counter disease while also mitigating host damage. Host TFs can modulate innate immune gene expression through diverse processes including chromatin remodeling, modification of transcriptional regulators, recruitment of coactivators, engagement of the Mediator complex, and recruitment of RNA Polymerase II (RNA Pol II) and its associated machinery to the transcriptional start site (TSS) ^1–4^. This increased presence of RNA Pol II ultimately promotes higher rates of initiation and elongation to drive expression of the gene target. *Cis*-regulatory elements such as promoters and enhancers act as an added layer of gene expression control for complex signaling pathways by providing unique DNA sites for TFs to bind and assemble ^5^. These enhancer sites can contain multiple motifs for distinct TF combinations which can alter chromatin architecture, recruit RNA Pol II, and initiate transcription in response to specific stimuli. These complexes termed ‘enhanceosomes’ are essential for regulation of the *IFNB1* promoter, which requires recruitment of the ATF2/c-Jun, IRF3/7, and NFκB TFs to sufficiently activate antiviral programing ^6,7^. As these TFs and regulatory elements are the ultimate mediators of antiviral responses, the molecular mechanisms underlying their assembly are of fundamental interest to the field of immunology.

Coordination of gene expression is complex, leading to the evolution of different classes of multifunctional factors that can bridge these protein-nucleic acid and protein-protein interactions. The Drosophila behavior/human splicing (DBHS) family are a trio of molecular scaffolds found in vertebrates and invertebrates that broadly associate with DNA, RNA, and TFs to mediate transcriptional repression and activation, initiation and elongation, and termination. The three DBHS members: splicing factor proline/glutamine rich (SFPQ/PSF), non-POU domain-containing octamer-binding protein (NONO/p54nrb) and paraspeckle protein component 1 (PSPC1/PSP1) are involved in every aspect of gene regulation including transcription, pre-mRNA splicing, RNA transport, and DNA repair. Transcriptional activation by DBHS proteins is driven by interactions with the transcriptional machinery, association with gene promoters, and processing of nascent RNA transcripts. These variety of functions demonstrate the utility of DBHS proteins as “molecular scaffolds” to facilitate a variety of activities related to gene regulation. However, their role in regulation of innate immune activation is unclear.

The role of DBHS proteins during viral infection has been controversial with investigations identifying both proviral and antiviral activities. This may be because DBHS proteins can regulate virus replication directly confounding effects on the host response. In particular, NONO has been shown to interact with the genomes for a variety of viruses to support their replication ^8–11^ while also functioning as an innate immune sensor for the HIV capsid ^12^. Given the diversity of DBHS protein functions, and the importance of paraspeckle components in viral replication and antiviral gene expression, we examined the antiviral capacity of NONO and revealed its role as a novel transcriptional regulator that broadly promotes innate immunity. NONO was required to elicit potent antiviral responses against a panel of orthoflaviviruses and innate immune agonists. NONO’s antiviral activity was mediated specifically at the transcriptional level as NONO enhances chromatin accessibility of innate immune genes. This leads to enhanced RNA Pol II recruitment and binding to promoter regions ultimately driving expression of antiviral genes. This work identifies an unrecognized regulatory role for NONO in antiviral innate immunity promoting efficient transcription of IFN-I, ISGs, and proinflammatory cytokines.

## Results

### NONO is required for establishing an effective antiviral response

We generated *NONO* knockout (KO) A549 cells using CRISPR/Cas9 (Figures S1A and S1B) and assessed whether loss of endogenous NONO impacted replication of a panel of orthoflaviviruses. Loss of NONO increased replication of several mosquito- and tick-borne orthoflaviviruses including West Nile virus (WNV) Bird 114 and NY99 strains, yellow fever virus (YFV) 17D and Asibi strains, Zika virus (ZIKV) Dakar, Dengue virus serotype 2 (DENV-2) NGC, and several related viruses in the tick-borne encephalitis virus (TBEV) serogroup: TBEV Sofjin, Omsk hemorrhagic fever virus (OHFV) Guriev, and Kyasanur forest disease virus (KFDV) P9605 (Figures 1A and S1C). The impact of NONO was most consistently observed late in multi-step growth curves, resulting in a 10-fold increase in titers of infectious virus for most of the strains tested by 72 hours post infection (hpi) (WNV, YFV-17D, TBEV, OHFV, DENV-2, and ZIKV) and a 5-fold increase for KFDV and YFV-Asibi (Figures 1A and S1C). Increased viral replication in the *NONO* KO cells was also demonstrated by higher accumulation of positive-strand viral RNA (vRNA) and protein expression for several of the orthoflaviviruses tested (Figures S1D-G).

**Figure 1.**
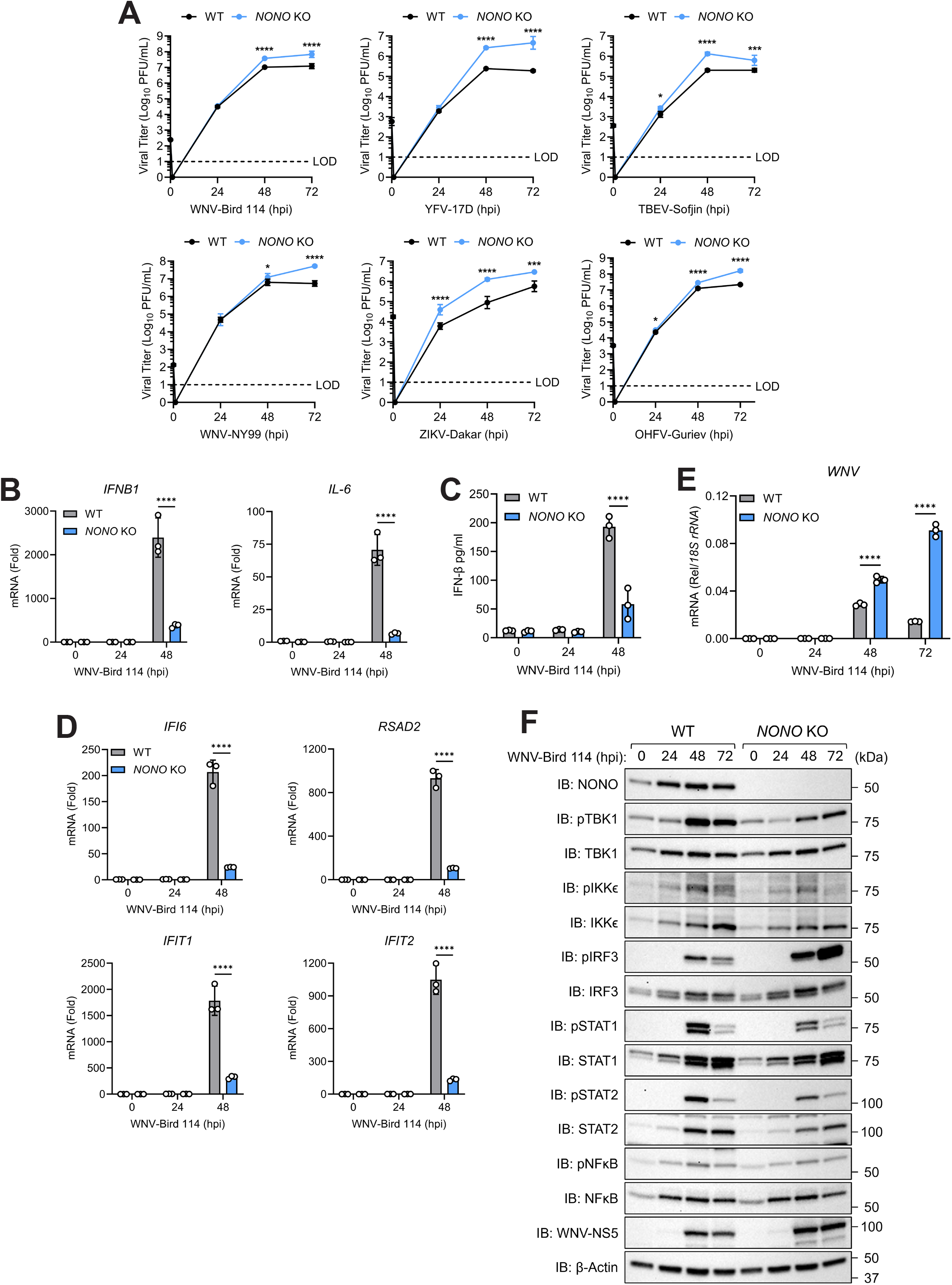
NONO loss enhances orthoflavivirus replication. (A) Infectious viral titers from WT or *NONO* KO A549 cells infected with West Nile virus (WNV) Bird 114 or NY99 MOI=0.0001, yellow fever virus (YFV) 17D MOI=0.001, Zika virus (ZIKV) Dakar MOI=0.01, tick-borne encephalitis virus (TBEV) Sofjin MOI=0.001, or Omsk hemorrhagic fever virus (OHFV) Guriev MOI=0.01 (plaque assay). (B-F) WT or *NONO* KO A549 cells infected with WNV-Bird 114 MOI=0.0001. (B) *IFN-β* and *IL-6* expression (qRT-PCR). (C) IFN-β protein in supernatants (ELISA). (D) *IFI6*, *RSAD2*, *IFIT1*, and *IFIT2* expression (qRT-PCR). (E) *WNV* expression (qRT-PCR). (F) Levels of WNV-NS5 protein and phosphorylation of TBK1, IKKε, IRF3, NFκB, STAT1, and STAT2 (immunoblot). Data are expressed as means (n=3) ± SD, *p < 0.05, ***p < 0.001, ****p < 0.0001 (two-way ANOVA with Sidak’s multiple comparisons) and are representative of 2-3 independent experiments.

We next examined whether NONO is associated with activation of antiviral innate immune responses. Expression of *IFNB1* and *IL-6* mRNAs were impaired in WNV-infected *NONO* KO cells compared to WT controls (Figure 1B and S1H). This correlated with lower levels of biologically active IFN-β measured in cell culture supernatants (Figure 1C and S1I) as well as lower mRNA expression for several IFN-stimulated genes (ISGs) shown to have antiviral activity against orthoflaviviruses (Figure 1D and S1J). In addition, *NONO* KO cells had reduced phosphorylation of TBK1, NFκB, STAT1, and STAT2 and an increase in WNV vRNA (Figure 1E and 1F) and NS5 protein (Figure S1K and S1L) consistent with reduced IFN signaling. However, higher levels of phosphorylated IRF3 were observed. This apparent disconnect between IRF3 phosphorylation and *IFNB1* transcription suggested NONO’s involvement in antiviral innate immunity occurs downstream of cytoplasmic signaling components.

Since loss of NONO was associated with a defective antiviral state, we next asked whether the increased viral replication observed in the *NONO* KO cells was due to the loss in IFN-I and ISG expression. To block innate immune signaling, WT and *NONO* KO cells were treated with the inhibitor Ruxolitinib which prevents JAK phosphorylation and subsequent downstream transcription of ISGs ^13,14^. Suppression of IFN signaling increased WNV and ZIKV replication in WT cells to levels equivalent to those observed in *NONO* KO cells (Figures 2A, 2B, and S2), but the presence of Ruxolitinib did not have an added effect on viral replication in the *NONO* KO cells. Infection with WNV induced higher *RSAD2*, *IFI6*, and *OAS1* mRNA levels in WT cells compared to *NONO* KO cells (Figure 2C) and treatment with Ruxolitinib prevented mRNA induction for these ISGs in both lines indicating successful inhibition of IFN signaling (Figure 2C). Additional confirmation by immunoblot revealed the DMSO-treated *NONO* KO cells have less phosphorylated STAT1 and STAT2 while also having higher levels of WNV NS5 compared to the WT (Figure 2D). Ruxolitinib treatment abolished STAT phosphorylation in both lines and equalized WNV NS5 expression in WT and *NONO* KO cells (Figure 2D). Taken together, these results indicate NONO is required for the optimal expression of antiviral innate immune genes and establishment of an antiviral state following infection with medically relevant orthoflaviviruses.

**Figure 2.**
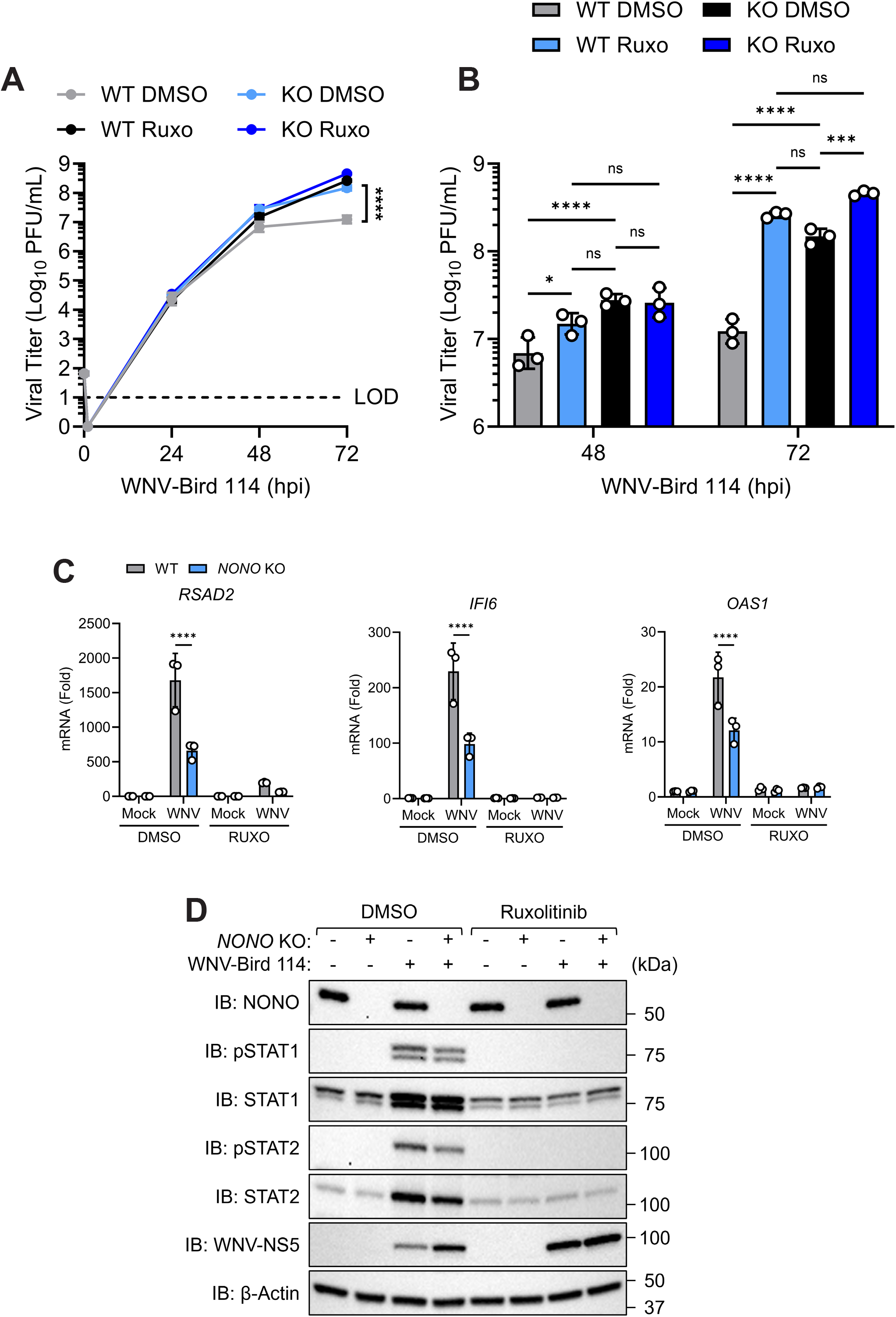
NONO is critically required for establishing an effective antiviral response. (A-D) WT or *NONO* KO A549 cells pretreated with Ruxolitinib (10 μg/mL) or DMSO carrier for 1 hour before infection with West Nile virus (WNV) Bird 114 MOI=0.0001. (A) Infectious viral titers (plaque assay). (B) Zoom in of 48-72 hpi infectious viral titers from (A). (C) *RSAD2*, *IFI6*, and *OAS1* expression 48 hpi (qRT-PCR). (D) Levels of WNV-NS5 protein and phosphorylation of STAT1 and STAT2 48 hpi (immunoblot). Data are expressed as means (n=3) ± SD, *p < 0.05, ***p < 0.001, ****p < 0.0001 (two-way ANOVA with Tukey’s or Sidak’s multiple comparisons) and are representative of 2-3 independent experiments.

### NONO broadly promotes innate immune responses

To determine if a role for NONO in innate immunity is specific to orthoflaviviruses, we examined the differential gene expression profiles between WT and *NONO* KO cells following SeV stimulation using RNA-seq. Loss of NONO led to a significant reduction in transcript abundance as well as Reads Per Kilobase Million (RPKM) for *IFNB1* in response to SeV infection, consistent with our initial findings (Figure S3A). Downregulated genes included multiple ISGs (*DDX58*, *IFITM1*, and *MX1*) (Figure 3A). Gene Set Enrichment Analysis (GSEA) ^15,16^ revealed downregulation of pathways involved in antiviral defenses including IFN-α/β signaling, IFN-γ signaling, and anti-inflammatory responses to SeV infection (Figure 3B). These specific pathways were not different in unstimulated WT and *NONO* KO cells suggesting a role for NONO following stimulation and not at basal levels (Figure 3C). Upregulated genes in both mock and SeV-infected *NONO* KO cells were enriched in pathways for prefoldin-mediated transfer of substrates to the T-complex protein ring complex/chaperonin containing TCP-1 (TRiC/CCT), cooperation of prefoldin and TRiC/CCT in actin and tubulin folding, DNA damage recognition in global genome nucleotide excision repair (GG-NER), and RNA Pol III transcription initiation (Figure S3B and S3C). Interestingly, the most negative GSEA changes in both mock and SeV-infected *NONO* KO cells were found in metabolic and calcium-dependent signaling pathways including calmodulin induced events, glucagon signaling in metabolic regulation, and GLP-1 regulated insulin secretion (Figures 3B and 3C). Calcium signaling has emerged as a crucial regulator of innate immunity particularly in airways ^17,18^ and NONO loss has been shown to negatively impact calmodulin levels ^19^. Glucose homeostasis and metabolic regulation are also crucial for activating innate immunity ^20,21^. NONO post-transcriptionally regulates mRNA expression of metabolic genes and NONO-deficient mice exhibit impaired glucose tolerance ^22^. Thus, NONO is integral to controlling antiviral responses in addition to supporting its multiple biological activities. Despite the general alterations to gene expression following *NONO* KO, only the innate response was impactful to WNV replication (Figures 2A and 2B) demonstrating a critical role for NONO in innate immunity.

**Figure 3.**
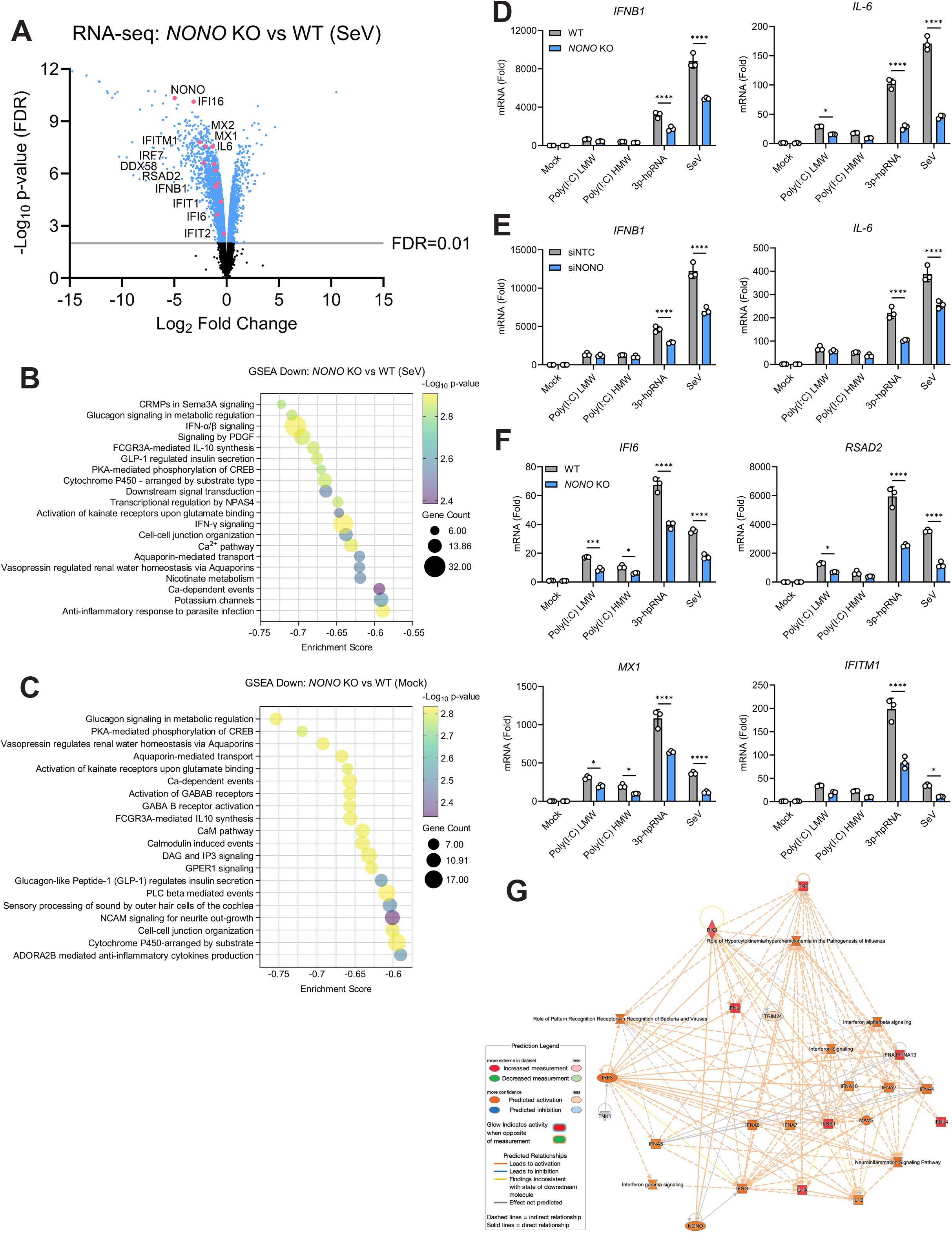
NONO broadly promotes innate immune responses. (A-C) WT or *NONO* KO A549 cells infected with SeV (100 HAU/mL) for 8 hours. (A) RNA-seq volcano plot of innate immune genes significantly downregulated in *NONO* KO compared to WT samples. (B-C) GSEA terms and pathways of downregulated transcripts from SeV-infected (B) and mock (C) samples. (D) *IFN-β* and *IL-6* expression from WT or *NONO* KO A549 cells stimulated with various PAMPs (1 μg/mL) or SeV (100 HAU/mL) for 8 hours (qRT-PCR). (E) *IFN-β* and *IL-6* expression from A549s treated with siNTC or siNONO and stimulated with various PAMPs (1 μg/mL) or SeV (100 HAU/mL) for 8 hours (qRT-PCR). (F) *IFI6, RSAD2, MX1,* and *IFITM1* expression from WT or *NONO* KO A549 cells stimulated with various PAMPs (1 μg/mL) or SeV (100 HAU/mL) for 8 hours (qRT-PCR). (G) A custom interaction network using the Ingenuity Pathway Analysis (IPA) Molecule Activity Predictor (MAP) tool visualizes predicted directional consequences of NONO activity on connected molecules, including *IFNB1*, and IFN family cytokines. Data are expressed as means (n=3) ± SD, *p < 0.05, ***p < 0.001, ****p < 0.0001 (two-way ANOVA with Sidak’s multiple comparisons) and are representative of 2-3 independent experiments.

Having established that NONO regulates the innate immune response to infection with RNA viruses, we next tested whether a role for NONO could also be observed with synthetic immune agonists. WT and *NONO* KO cells were treated with pathogen associated molecular patterns (PAMPs) known to stimulate specific pattern recognition receptors (PRRs). Transcript levels for *IFNB1*, *IL-6*, and several ISGs were significantly reduced in *NONO* KO cells compared to WT 8 hours after stimulation with SeV and 3p-hpRNA, an *in vitro* transcribed hairpin RNA of NEP from segment 8 of influenza A virus specifically recognized by RIG-I ^23,24^ (Figure 3D-F). Additionally, these results were replicated using a pool of small interfering RNAs (siRNA) to deplete endogenous NONO (Figures 3E, S3D, and S3E). Stimulating *NONO* KO cells with either low or high molecular weight Poly(I:C), dsRNA mimetics that stimulate RIG-I or MDA5 respectively, modestly impaired *IFNB1*, *IL-6*, and ISGs compared to WT at this early timepoint (Figure 3D-F). Furthermore, a kinetics study using Poly(dA:dT), a dsDNA mimetic sensed by several PRR sensors, identified a defect in *IFNB1*, *IL-6*, and *IFI6* transcription in *NONO* KOs beginning at 4 hours post stimulation (Figure S3F). *NONO* KO cells displayed an inability to mount an effective antiviral response to Poly(dA:dT) with increased transcriptional deficit over time (Figure S3F). To simulate the directional consequences of NONO activity on antiviral molecules, an Ingenuity Pathway Analysis (IPA) network comparing mock to SeV-infected WT cells was generated ^25^ (Figure S3G) and a custom interaction network using the Molecule Activity Predictor (MAP) tool was constructed in which NONO was manually added. The IPA MAP visual analysis predicted NONO as having an activating relationship for *IFNB1* and IFN family cytokines (Figure 3G). These results identify NONO as a regulatory factor that is essential for expression of IFN-I and ISGs following a wide range of innate immune stimuli.

### NONO augments innate immunity specifically at the transcriptional level

To assess whether NONO functions specifically at the transcriptional level, we independently evaluated signaling factor activation and gene transcription in both the IFN production and signaling pathways. Stimulation of the IFN production pathway with SeV in *NONO* KO cells replicated the increase in phosphorylation for IRF3 observed during WNV infection (Figure 4A). Consistent with our prior experiment, this higher activation of IRF3 was divorced from the lower *IFNB1* transcription and translation seen in the *NONO* KO. Expression of *IFNB1* was reduced in *NONO* KO cells beginning at 4 hpi leading to lower overall levels of IFN-β protein expression (Figures 4B and 4C). As expected, this diminished induction of IFN-I was associated with lower transcription to numerous ISGs across the time course (*IFI6*, *RSAD2*, *MX1*, and *IFIT1*) (Figure 4D). To determine whether this loss of ISG expression is direct or because of reduced expression of *IFNB1*, cells were stimulated with exogenous IFN-β. Phosphorylation of STAT1 or STAT2 after direct IFN-β stimulation was equivalent in WT and NONO KO cells (Figure 4E), yet loss of NONO resulted in a 5-fold lower induction of *IFI6*, *RSAD2*, and *IFIT1* suggesting that NONO has the potential to impact amplification of the IFN-I response through ISG transcription (Figure 4F). NONO transcript levels were not increased in response to SeV infection or IFN-β treatment (Figure S4A and S4B), demonstrating NONO is not an ISG under these conditions. Subcellular fractionation of cells infected with SeV or WNV confirmed that NONO itself does not relocalize from its nuclear compartment to the cytoplasm indicating NONO is unlikely to directly regulate cytoplasmic signaling events (Figures 4G and S4C). In support of this, there was no defect in the cytoplasmic-nuclear translocation of activated innate immune transcription factors (phosphorylated IRF3, STAT1, and NFκB) in SeV-infected *NONO* KO cells (Figure 4G). Furthermore, we did not observe endogenous NONO binding with endogenous STAT1 or STAT2 upon IFN-β stimulation. We also did not detect enhanced endogenous binding between NONO and its other DBHS partners under these conditions (Figure S4D).

**Figure 4.**
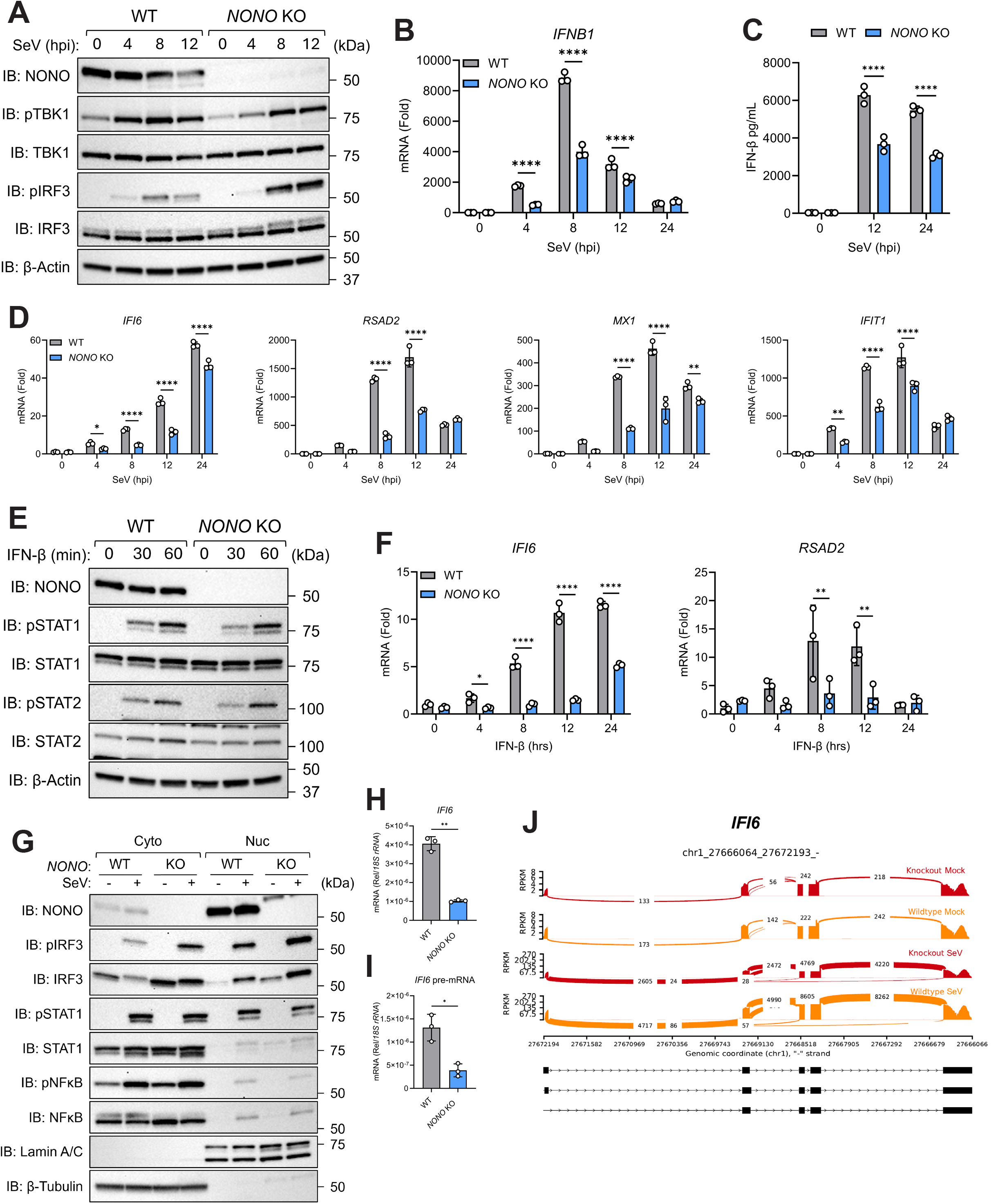
NONO augments innate immunity specifically at the transcriptional level. (A-D) WT or *NONO* KO A549 cells infected with SeV (100 HAU/mL). (A) Phosphorylation levels of TBK1 and IRF3 (immunoblot). (B) *IFN-β* expression (qRT-PCR). (C) IFN-β protein in supernatants (ELISA). (D) *IFI6, RSAD2, MX1,* and *IFIT1* expression (qRT-PCR). (E) Phosphorylation levels of STAT1 and STAT2 from WT or *NONO* KO A549 cells stimulated with IFN-β (1000 IU/mL) (immunoblot). (F) *IFI6* and *RSAD2* expression from WT or *NONO* KO A549 cells stimulated with IFN-β (100 IU/mL) (qRT-PCR). (G) Subcellular fractionation of WT or *NONO* KO A549 cells infected with SeV (100 HAU/mL) for 8 hours (immunoblot). (H) Mature and pre-mRNA expression for *IFI6* from WT or *NONO* KO A549 cells stimulated with IFN-β (100 IU/mL) for 8 hours (qRT- PCR). (I) Sashimi plots for *IFI6* from WT or *NONO* KO A549 cells mock or SeV infected (100 HAU/mL) for 8 hours. Data are expressed as means (n=3) ± SD, *p < 0.05, **p < 0.01, ****p < 0.0001 (two-way ANOVA with Sidak’s multiple comparisons or Student’s t-test) and are representative of 2-3 independent experiments.

NONO engages in nearly every step of gene regulation including post-transcriptional regulation of gene expression by binding pre-mRNAs to enhance splicing ^26–29^. To determine whether NONO promotes ISG induction by regulating splicing of ISGs, we performed replicate multivariate analysis of transcript splicing (rMATS) on our RNA-seq data, which quantifies alternative splicing events and identifies differentially regulated exons across samples ^30^. This analysis revealed that most of the significant alternative splicing events in *NONO* KO cells include skipped exons and retained introns, indicating that a dysregulation of splicing does occur (Figure S4E). However, when comparing levels of mRNA and pre-mRNA for *IFI6* and *MX1* as example ISGs between WT and *NONO* KOs, accumulation of the pre-mRNA form for either of these ISGs was not observed (Figures 4H, 4I, S4F, and S4G). We also did not detect significant changes in alternative splicing events for mRNA isoforms of *IFI6*, *MX1*, *RSAD2*, and *IL-6* (Figures 4J, and S4H-J). Additionally, none of the differentially expressed ISGs in *NONO* KOs were ranked as having significantly altered splicing events suggesting NONO’s function in antiviral innate immunity is independent of its role as a splicing factor (Figure S4K). Collectively, these data identify NONO as a critical component for innate immunity and strongly suggests NONO functions as a transcriptional regulator of IFN-I and ISG expression.

### NONO enhances chromatin accessibility of innate immune genes

Recently, NONO was found to be integral for proper H3K36me2 active chromatin formation and neural differentiation of stem cells ^31–33^. To test the hypothesis that NONO-dependent chromatin alterations are also required for efficient activation of antiviral innate immunity, we implemented ATAC-seq to assess genome-wide chromatin accessibility sites in WT and *NONO* KO cells. Upon quantification of ATAC-seq signal, we observed a slight increase in chromatin accessibility in *NONO* KO cells compared to WT under basal conditions (Figure 5A). These differences in accessibility were erased after SeV infection indicating loss of NONO does not broadly affect chromatin remodeling (Figure 5B). Principal component analysis identified a mild distinction between genotypes (18% for WT to *NONO* KO comparisons) compared to the more apparent differences attributed to treatment (73% for mock to SeV comparisons) (Figure 5C), supporting the conclusion that NONO loss does not lead to substantial remodeling of the chromatin environment across the transcriptome. However, the chromatin sites that were significantly reduced in the infected *NONO* KOs corresponded to innate immune genes that were also differentially expressed in our RNA-seq analysis (*IFNB1*, *RSAD2*, *MX1*, *DDX58*, *IFIT1*, *IFIT2* and *IL-6*) (Figure 5D). Similarly, Integrative Genomics Viewer tracks depicting the genomic regions containing *IFNB1*, *IL-6* and its lncRNA enhancer *IL6-AS1*, *RSAD2*, and *CMPK2* identify regions with reduced chromatin accessibility in the *NONO* KO during SeV infection (Figures 5E, S5A, and S5B).

**Figure 5.**
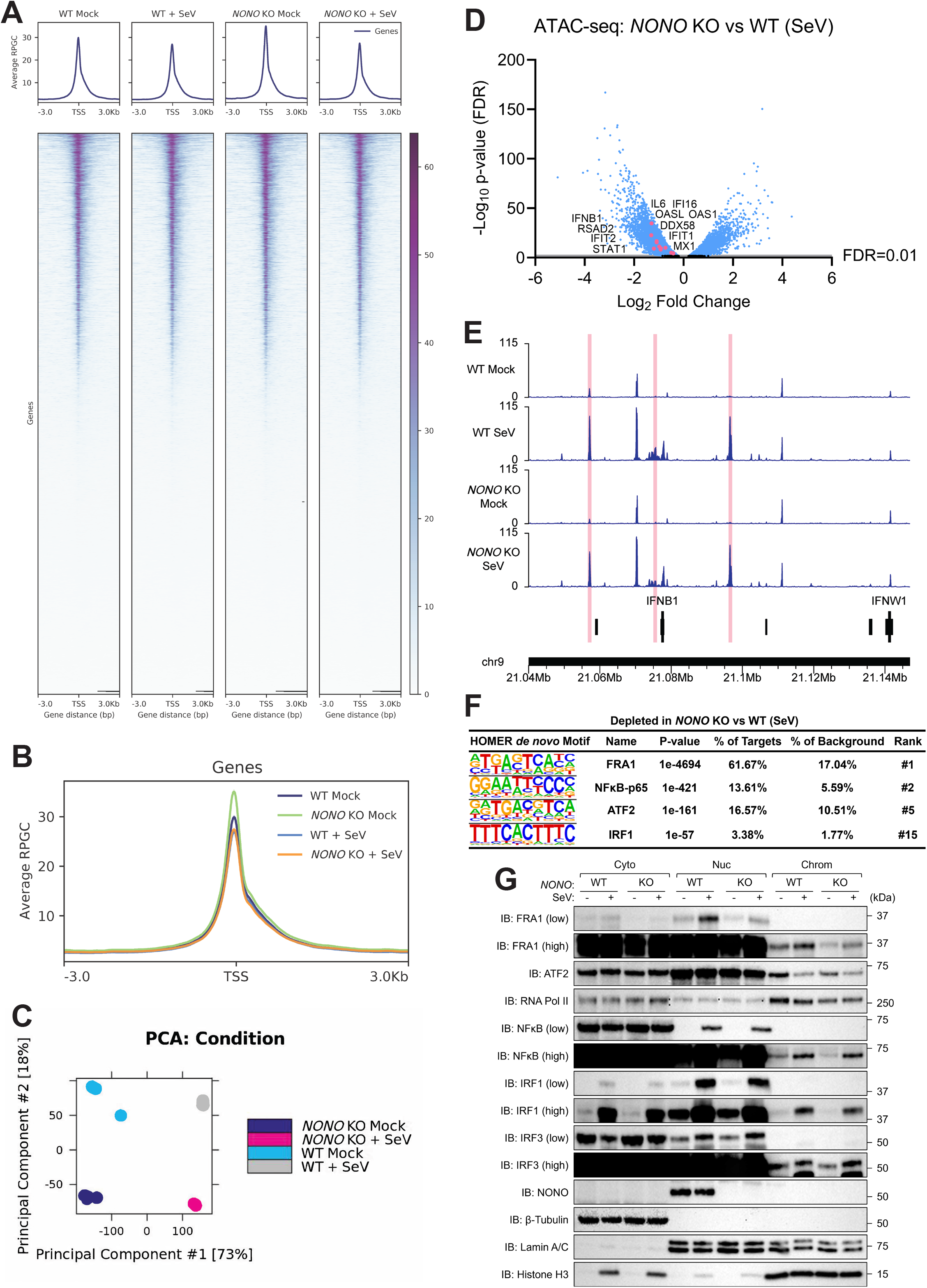
NONO enhances chromatin accessibility of innate immune genes. (A-F) ATAC-seq of WT or *NONO* KO A549 cells infected with SeV (100 HAU/mL) for 8 hours (n=4 per group). (A) ATAC-seq signal for genome-wide chromatin accessibility (signal shown ± 3 Kb from TSS). (B) Average ATAC-seq profile plots for chromatin accessibility (signal shown ± 3 Kb from TSS). (C) Multidimensional PCA plot of ATAC-seq samples demonstrate a large effect of treatment (component #1), and a mild effect of genotype (component #2). (D) ATAC-seq volcano plot for chromatin peaks of innate immune genes significantly less accessible in SeV-infected *NONO* KO compared to WT samples. (E) Genome browser view of open chromatin peaks for *IFNB1* with highlighted portions indicating regions with significantly less available chromatin in the SeV-infected *NONO* KO compared to the SeV-infected WT. (F) HOMER *de novo* motif analysis for depleted motifs in SeV-infected *NONO* KO compared to WT samples. (G) Subcellular fractionation of WT or *NONO* KO A549 cells infected with SeV (100 HAU/mL) for 8 hours (immunoblot).

We next used motif discovery analysis to identify transcription factor binding sites where chromatin accessibility was impacted by NONO deletion. HOMER *de novo* motif discovery identified sites with depleted accessibility in *NONO* KO cells to be strongly enriched for motifs corresponding to components of the AP-1 complex (FOS and ATF2) under basal conditions (Figure S5C). The availability of AP-1 motifs (FRA1 and ATF2) as well as others relevant for innate immune transcription factor binding (NFκB-p65 and IRF1) were strongly reduced in SeV-stimulated *NONO* KO cells (Figure 5F). In agreement with these findings, subcellular fractionation revealed FRA1, ATF2, and RNA Pol II to have a reduced presence in the chromatin-bound compartment for *NONO* KO cells under basal conditions (Figure 5G). However, chromatin binding of other TFs including NFκB, IRF1, and IRF3 were unaffected demonstrating specificity. The decrease in FRA1 and RNA Pol II could also be observed when comparing chromatin-bound levels between SeV-infected *NONO* KO and WT cells. Together, our ATAC-seq and motif analysis data indicate that loss of NONO results in reduced chromatin accessibility for antiviral genes and diminished innate immune transcription factor motif availability at DNA binding sites leading to lower association levels of the RNA Pol II transcriptional machinery.

### NONO promotes RNA Pol II binding to the promoter regions of innate immune genes

To verify that reduced chromatin accessibility and transcription factor binding contributed to the impairment in antiviral gene transcription in *NONO* KO cells, we performed chromatin immunoprecipitation (ChIP) targeting RNA Pol II followed by qRT-PCR to examine binding of the transcriptional machinery to promoter regions of innate immune genes. ChIP-qPCR revealed *NONO* KO cells had reduced RNA Pol II binding to the promoter region for *IFNB1* during SeV infection (Figure 6A). Several factors may alter transcription factor binding preferences, including dynamic changes in chromatin state. We focused on chromatin structure based on our ATAC-seq results implicating a role for NONO in chromatin availability for innate immune genes. Histone modifications like methylation can be added to the lysine residues of H3 histones and are implicated in both transcriptional activation and repression depending on the methylation site ^34^. These changes can result in rapid chromatin remodeling altering gene regulation in response to stimuli. We performed ChIP-qPCR on WT and *NONO* KO cells using antibodies for either an open, transcriptionally active chromatin state (H3K4me3) or a closed, transcriptionally repressed form (H3K27me3) following SeV infection. H3K4me3 was strongly associated with the *IFNB1* promoter in SeV-infected WT cells compared to the *NONO* KO (Figure 6B). Conversely, H3K27me3 levels on the *IFNB1* promoter were reduced in WT cells while remaining steady in the *NONO* KO after infection (Figure 6B).

**Figure 6.**
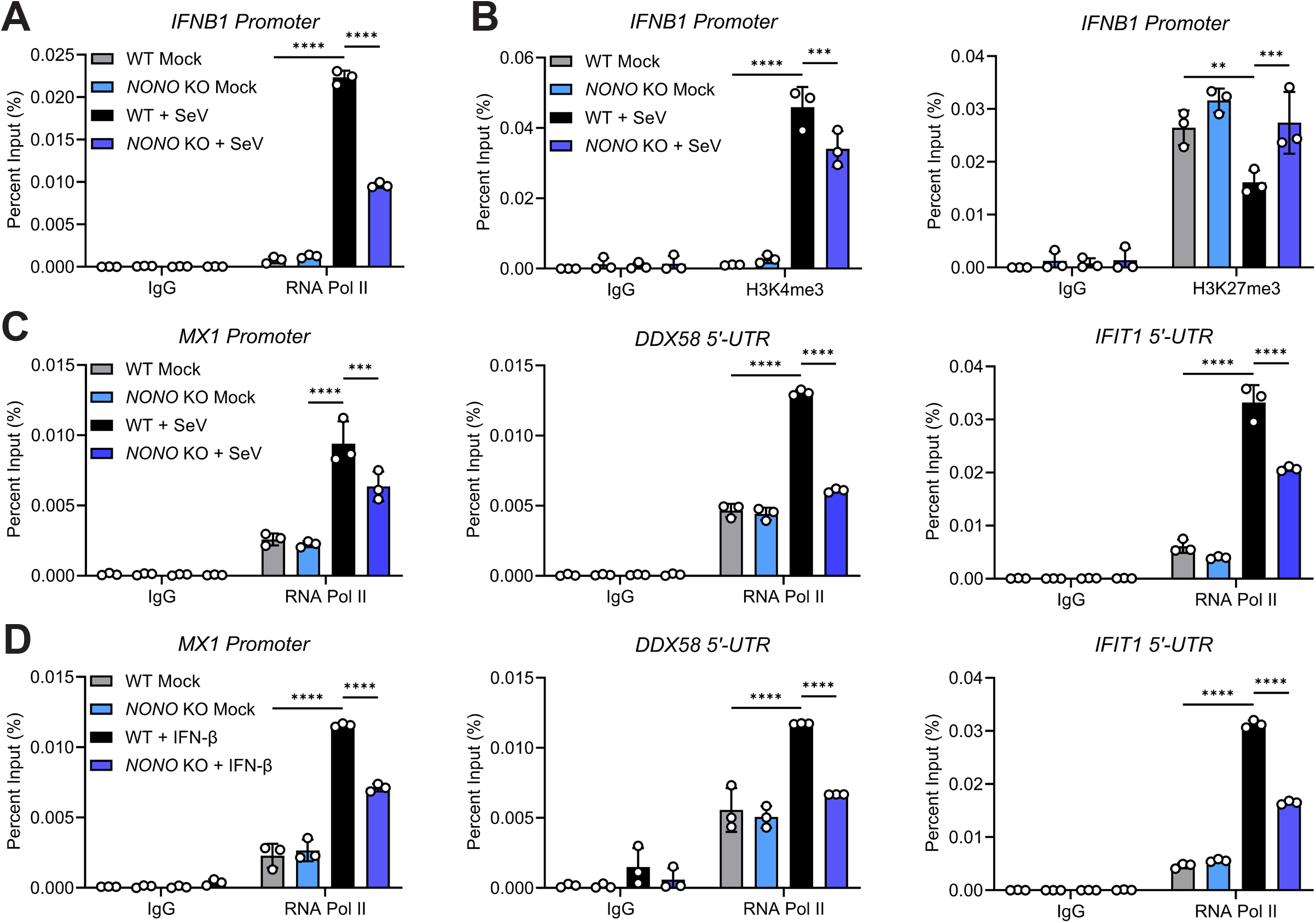
NONO loss impairs RNA Pol II binding to innate immune gene promoters. (A-C) WT or *NONO* KO A549 cells infected with SeV (100 HAU/mL) for 8 hours. (A) ChIP-qPCR showing RNA- Pol II occupancy at the *IFN-β* promoter. (B) ChIP-qPCR showing either H3K4me3 or H3K27me3 histone modifications at the *IFN-β* promoter. (C) ChIP-qPCR showing RNA-Pol II occupancy at the *MX1* promoter, *DDX58* 5’-UTR, or *IFIT1* 5’-UTR. (D) ChIP-qPCR showing RNA-Pol II occupancy at the *MX1* promoter, *DDX58* 5’-UTR, or *IFIT1* 5’-UTR for WT or *NONO* KO A549 cells stimulated with IFN-β (1000 IU/mL) for 8 hours. Data are expressed as means (n=3) ± SD, **p < 0.01, ***p < 0.001, ****p < 0.0001 (two-way ANOVA with Tukey’s multiple comparisons) and are representative of 2-3 independent experiments.

We next examined binding of RNA Pol II to the promoters or 5’ untranslated regions (5’-UTR) by ChIP-qPCR for three ISGs that were differentially expressed in the SeV-infected *NONO* KO cells (*MX1*, *DDX58*, and *IFIT1*). RNA Pol II was associated with each of the three ISG targets in the SeV-infected WT cells compared to the mock. In contrast, transcriptional machinery binding to these sites was significantly impaired in *NONO* KO cells (Figure 6C). Furthermore, direct stimulation of the IFN-I signaling pathway with IFN-β enhanced RNA Pol II association with these ISG promoters while reproducing the observed binding deficit in the absence of *NONO* (Figure 6D). These effects of RNA Pol II binding in response to NONO loss were not observed for unrelated control genes including RPL30 and the α satellite repeat element (Figure S6A-C). The RPL30 gene is actively transcribed in all cell types, and its promoter is highly enriched for H3K4me3 histone modifications while the α satellite repeat element found in pericentromeric regions of chromosomes shows hallmarks of transcriptionally silent heterochromatin. Taken together, the data demonstrates that NONO impacts chromatin architecture, access for transcriptional machinery, and ultimately gene expression for numerous antiviral innate immune genes.

## Discussion

In this study, we identify NONO as a host factor required for the restriction of several orthoflaviviruses through the transcription of IFN-I and ISGs necessary for establishing an antiviral state. These findings reveal a role for NONO as an enhancer of innate immune induction as loss of NONO significantly reduced levels of IFN-I and ISGs through direct impairment of transcription. This function of NONO was observed following stimulation with synthetic analog PAMPs suggesting NONO may function as a broader regulator of antiviral and pro-inflammatory programs. A mechanistic dissection of gene transcription demonstrated that NONO is required for chromatin accessibility of multiple innate immune genes to enable RNA Pol II binding to their promoter regions. Taken together, our work greatly extends the current understanding of NONO in gene regulation by establishing NONO as a key transcriptional component for broad antiviral responses and by discerning a mechanism for how NONO potentiates transcriptional machinery’s induction of antiviral innate immunity.

The viruses whose replication was impacted by NONO’s function in innate immune transcription included WNV, YFV-17D, ZIKV, TBEV, OHFV, DENV-2, and KFDV representing diverse members of the orthoflavivirus genus with varying degrees of pathogenicity. Interestingly, loss of NONO differentially enhanced orthoflavivirus replication, even across strains of the same virus. YFV-17D displayed a 10-fold increase in titers compared to the 5-fold increase for YFV-Asibi in *NONO* KO cells. YFV-17D is highly sensitive to IFN-I and elicits a stronger immune response compared to the more virulent Asibi strain ^35^ which may explain why some viruses displayed improved replication in the *NONO* KO cells. Orthoflaviviruses evade the host innate immune response due to shielding of their viral replication intermediates within remodeled membranes of the endoplasmic reticulum in addition to specific virus-encoded mechanisms that antagonize and delay IFN-I expression by infected cells ^36–38^. However, the effects of NONO on IFN-I and ISGs were evident as early as 4 to 8 hours post stimuli by defined agonists (PAMPs, IFN-β, and SeV) suggesting that NONO is required for early gene expression. NONO is predominantly a nuclear protein capable of forming paraspeckles and stress granules under different conditions ^39^, yet WNV and SeV infection did not lead to a shift in NONO levels between cytoplasmic and nuclear compartments as seen with another DBHS member, SFPQ, during EMCV infection ^40,41^. NONO’s function in IFN-I and ISG expression was at the transcriptional level and independent of cytoplasmic signaling events despite the observed increase in IRF3 phosphorylation in *NONO* KOs. Previous studies indicated that swine NONO promotes antiviral innate immune responses by acting as a positive regulator of swine IRF3 to inhibit porcine reproductive and respiratory syndrome virus ^42,43^. In contrast, human NONO did not impair the phosphorylation or cytoplasmic-nuclear translocation of endogenous human IRF3, NFκB, or STAT1. It is not immediately apparent why IRF3 phosphorylation at S396 was enhanced in the *NONO* KO cells since protein and transcript abundance for upstream kinases TBK1 and IKKε remained unchanged. One explanation could be that NONO is involved in gene regulation for a protein phosphatase that regulates IRF3 activity. Although we did not detect significant changes in the expression of the known IRF3 phosphatase PTPA or its adaptor RACK1 ^44^ by RNA-seq, the possibility remains that NONO is important for the expression of a yet uncharacterized IRF3 phosphatase.

Reports have identified proviral roles for DBHS paraspeckle proteins suggesting these three host factors directly and indirectly assist viral replication ^41^. SFPQ is a known target of Encephalomyocarditis virus (EMCV) and Coxsackievirus B3 (CVB3) acting as an internal ribosome entry site (IRES) trans-acting factor (ITAF) directly promoting vRNA translation ^40,45^. SFPQ’s nuclear localization was altered during EMCV and CVB3 infection where it translocated to the cytoplasm to bind vRNA. However, no change in cellular distribution for SFPQ was observed following Poly(I:C) stimulation which was consistent with our findings for NONO during WNV or SeV infection. In another example, PSPC1 and NONO indirectly support herpes simplex virus 1 (HSV- 1) replication by recruiting the host transcription factor, STAT3, to paraspeckles. Once present, STAT3 interacts with promoters for viral lytic cycle genes like ICP0 to enhance activation and increase viral gene expression ^9^. Interestingly, SFPQ was found to negatively impact HSV-1 replication and knockdown of SFPQ alone increased ICP0 transcription ^9^. Structural data on DBHS proteins show they almost always operate as heterodimers with other DBHS members to execute their various functions ^26,27^. This heterodimer requirement may explain the frequency with which pairs of DBHS proteins are found to aid in viral replication.

A logical question that arises from these studies is why are DBHS proteins like NONO a target for numerous viruses? One explanation could be in how viruses utilize DBHS factors to fully augment their replication cycle. Viral genomes are limited in size and cannot encode a sufficient proteome to complete viral replication. Despite having a ∼30 Kb genome, SARS-CoV-2 still needs to overcome deficiencies in its genetic information by hijacking host RNA-binding proteins (RBPs) and repurposing them as their own machinery. Indeed, NONO and SFPQ have been identified in genome-wide RNA-interactome studies focused on host RBPs and SARS-CoV-2 vRNA ^46–49^. NONO can bind single and double-stranded RNA and DNA allowing it to interact with chromatin and pre-mRNA to greatly influence gene expression. This substrate promiscuity, coupled with the ability to form biomolecular condensates to mediate additional protein-protein interactions, make DBHS protein desirable targets for viruses. The evolutionary advantage of usurping DBHS proteins may be for viruses to gain additional means for vRNA transcription, splicing, and transport while also sequestering these host factors from participating in the antiviral response.

Collectively, our study identifies NONO as a restriction factor against several orthoflaviviruses and a critical component for optimal IFN-I and ISG transcription against a broad range of innate immune agonists. These findings provide new insights on the mechanisms by which DBHS proteins modulate host transcription by establishing NONO as an enhancer of chromatin accessibility for numerous antiviral genes. Developing a clearer understanding for how RBPs like NONO alter a gene’s epigenetic state will be important for unlocking new strategies in the development of future therapeutics for viral diseases and autoimmune disorders.

## Acknowledgments

This research was supported by the Intramural Research Program of the National Institutes of Health (NIH). The contributions of the NIH authors are considered Works of the United States Government. The findings and conclusions presented in this paper are those of the authors and do not necessarily reflect the views of the NIH or the U.S. Department of Health and Human Services.

## Author contributions

A.H. performed all aspects of this study. M.J. performed experiments. B.S., J.G.S., and R.M.B. provided critical reagents and technical advice. T.E.M., S.Y., P.A.B., J.B.L., and C.M. performed analysis of sequencing datasets. A.H. and S.M.B. organized the study and prepared the manuscript. All authors have read and agreed to the published version of the manuscript.

## Declaration of interests

The authors declare no conflicts of interest.

## RESOURCE AVAILABILITY

### Lead contact

Further information and requests for resources and reagents should be directed to and will be fulfilled by the lead contacts, Adam Hage and Sonja M. Best.

### Materials availability

Resources and reagents generated in this study are available upon request from the lead contacts.

### Data and code availability

Transcriptomic and epigenomic data generated during this study have been deposited to the NCBI Gene Expression Omnibus (GEO) database under GEO: GSE335258 (RNA-seq) and GEO: GSE335260 (ATAC-seq). All datasets are publicly available as of the date of publication.

This paper does not report original code.

Any additional information required to reanalyze the data reported in this study is available from the lead contacts upon request.

## EXPERIMENTAL MODEL AND SUBJECT DETAILS

### Cell lines

A549 (CCL-185) and Vero (CCL-81) cell lines were purchased from ATCC. All cells were maintained in Dulbecco’s Modified Eagle’s Medium (DMEM) (Gibco) supplemented with 10% (v/v) fetal bovine serum (FBS) (Gibco) and 1% (v/v) penicillin-streptomycin (Gibco). Cells used for transient transfections were plated in DMEM supplemented with 10% (v/v) FBS lacking 1% (v/v) penicillin-streptomycin.

### Viruses

Viruses used in this study were handled under biosafety level 2 (BSL-2), BSL-3, and BSL-4 conditions at the Rocky Mountain Laboratories Integrated Research Facility in accordance with Division of Select Agents and Toxins (DSAT) regulations for study of select agents and Institutional Biosafety approvals. Zika virus (Dakar), Yellow fever virus (Asibi), Tick-borne encephalitis virus (Sofjin), Kyasanur Forest disease virus (P9605), and Omsk hemorrhagic fever virus (Guriev) were kindly provided by The World Reference Center of Emerging Viruses and Arboviruses (WRCEVA) (The University of Texas Medical Branch at Galveston). West Nile virus (Bird 114 and NY99), Yellow fever virus (17D), and Dengue virus serotype 2 (NGC) were obtained from BEI Resources. Sendai virus (Cantell) was obtained from AVS Bio.

## METHOD DETAILS

### Transfections and stimulations

Transient transfection of PAMPs (Poly(I:C), 3p-hpRNA, and Poly(dA:dT) (InvivoGen)) for stimulations was performed by transfecting PAMPs with Lipofectamine 2000 (Invitrogen) in a 1 μg PAMP:1 μL Lipofectamine 2000 ratio prepared in Opti-MEM (Gibco) according to the manufacture’s guidelines.

### Cytokines and inhibitors

IFN-β stimulations were performed with human IFN-β 1a diluted in DMEM to a concentration of 100 or 1000 IU/mL (PBL Assay Science). IFN-I signaling blockade was performed with Ruxolitinib or DMSO control diluted in DMEM to a concentration of 10 μg/mL (InvivoGen). Cells were pretreated with Ruxolitinib or DMSO for 1 hour before infection and compounds were maintained at the indicated concentration in culture for the duration of the experiment.

### Immunoblot assay

Cell lysates were resolved on 10% or 8-16% Novex Tris-Glycine gels (Invitrogen) and 4-15% Criterion TGX gels (Bio-Rad) and transferred to polyvinylidene difluoride (PVDF) membranes using the Trans-Blot Turbo transfer system (Bio-Rad). Membranes were blocked with 5% (w/v) non-fat dry milk in TBST (TBS with 0.1% (v/v) Tween-20) for 1 hour, washed with TBST three times for 5 minutes each, and probed with the indicated primary antibody in 3% (w/v) BSA in TBST at 4°C overnight. Following overnight incubation, membranes were washed with TBST three times for 5 minutes each, probed with anti-rabbit or anti-mouse IgG (whole molecule)- peroxidase antibody produced in goat (Sigma) in 5% (w/v) non-fat dry milk in TBST for 1 hour at room temperature, and washed with TBST three times for 5 minutes each. Proteins were visualized by ECL (Thermo Scientific) or SuperSignal West Femto chemiluminescence reagents (Thermo Scientific) and detected using an iBright FL1500 Imaging System (Invitrogen). The primary antibodies and concentrations used are listed in the key resources table.

### Infections and plaque assays

A549 cells were seeded in 24-well plates (100,000 cells/well) and infected with virus diluted in DMEM at 37°C for 1 hour. Inoculations were removed and cells were washed once with DPBS, overlaid with DMEM containing 2% (v/v) FBS, 1% (v/v) penicillin streptomycin, and incubated at 37°C. Supernatants were collected for plaque assay at the indicated time points. For plaque assays, confluent monolayers of Vero cells were inoculated with supernatants serially diluted in DMEM containing 2% (v/v) FBS, 1% (v/v) penicillin streptomycin, and incubated at 37°C for 1 hour. Inoculums were removed and replaced with MEM containing 1.5% (w/v) carboxymethylcellulose and 1% (v/v) penicillin streptomycin and incubated at 37°C for 4 days (6 days for DENV-2 and 7 days for YFV). Cells were fixed in 10% (w/v) formalin for 1 hour at room temperature and stained with 1% (w/v) crystal violet for 10 minutes at room temperature.

### Quantitative reverse transcription PCR (qRT-PCR)

Total RNA isolation and gDNA removal was attained using the RNeasy Plus Mini Kit with gDNA Eliminator columns (Qiagen). cDNA synthesis was achieved using SuperScript VILO Master Mix (Invitrogen). qRT-PCR was performed using SsoAdvanced Universal SYBR Green Supermix (Bio-Rad) in a 384-well QuantStudio 7 Pro Real-Time PCR System (Applied Biosystems). Gene expression was normalized to human 18S rRNA by the comparative CT method (ΔΔCT). The primer sequences used are listed in Table S1.

### Chromatin immunoprecipitation (ChIP) and ChIP-qPCR

Chromatin was sheered using a Q800R3 Sonicator (Qsonica) at 70% amplitude for a total of 30 minutes on time using 15 second on and 45 second off intervals. ChIP was performed using the SimpleChIP Plus Sonication Chromatin IP Kit (Cell Signaling Technology) according to the manufacturer’s instructions. 2 μg of ChIP-grade antibodies were used for each immunoprecipitation. ChIP-qPCR was performed using SimpleChIP Universal qPCR Master Mix (Cell Signaling Technology) in a 384-well QuantStudio 7 Pro Real-Time PCR System (Applied Biosystems). Immunoprecipitation efficiency was calculated using the percent input method (2% x 2^(CT^ ^2%^ ^Input^ ^Sample-CT^ ^IP^ ^Sample)^). The primer sequences used are listed in Table S1.

### Co-immunoprecipitation

Cells were harvested in RIPA lysis buffer (50 mM Tris-HCl, pH 8.0, 150 mM NaCl, 1% (v/v) IGEPAL CA-630, 0.5% (w/v) sodium deoxycholate, 0.1% (v/v) SDS, and protease inhibitor cocktail (Roche). Cell lysates were clarified by centrifugation at 21,000 x RCF for 20 min at 4°C. 10% of the clarified lysate was added to 2X Laemmli buffer containing 2-Mercaptoethanol, heated for 10 min at 95°C, and stored at -20°C as whole-cell lysates (WCL). For endogenous proteins, lysates were subjected to immunoprecipitation with 2 μg of primary antibody overnight at 4°C followed by incubation with protein A/G agarose beads (Thermo Scientific) for two hours at 4°C on a rotating platform. Beads were washed seven times in RIPA buffer and bound proteins were eluted by heating samples in 2X Laemmli buffer containing 2-Mercaptoethanol for 10 min at 95°C.

### IFN-β ELISA

Secreted IFN-β was measured using the LumiKine Xpress hIFN-β 2.0 ELISA kit (InvivoGen) according to the manufacturer’s instructions.

### Cellular fractionation

Nuclear, cytoplasmic, and chromatin-bound compartments were separated using either the NE-PER nuclear and cytoplasmic extraction kit (Thermo Scientific) or the Subcellular Protein Fractionation Kit for Cultured Cells (Thermo Scientific) according to the manufacturer’s instructions.

### NONO siRNA knockdown

Transient knockdown of endogenous NONO in A549 cells, seeded in 24-well plates (30,000 cells per well), was achieved by transfection of ON-TARGETplus Non-targeting Control Pool (D-001810-10-05 Dharmacon) or SMARTpool: ON-TARGETplus NONO siRNA (L-007756-01-0005 Dharmacon) at a final concentration of 20 nM siRNA for 48 hours. Delivery of siRNA was achieved with Lipofectamine RNAiMAX (Invitrogen) according to the manufacture’s guidelines.

### Generation of NONO CRISPR knockout cells

A549 cells were electroporated using the SF Cell line 4D-Nucleofector X Kit S (Lonza) following the manufacturer’s recommended protocol. Briefly, 200,000 cells were resuspended in 20 μL of SF 4D- Nucleofector X solution containing 10 μg of Alt-R S.p. Cas9 Nuclease V3 (IDT) and 2 μL of 100 μM Alt-R CRISPR-Cas9 sgRNA (IDT) before electroporation using the CM-130 program. Cells were resuspended in DMEM and seeded in a 6-well plate before single-cell sorting into 96-well plates. Immunoblot was performed on cell lysates from individual monoclonal populations to identify *NONO* knockout cells. The sgRNA sequence used was 5’-TGGCAATCTCCGCTAGGGTT-3’.

### RNA-seq data processing

Total RNA isolation and gDNA removal was attained using the RNeasy Plus Mini Kit with gDNA Eliminator columns (Qiagen) according to the manufacturer’s instructions. RNA-seq libraries were prepared using the NEBNext Ultra II DNA Library Prep Kit according to the manufacturer’s instructions. RNA-seq libraries were sequenced on a NovaSeq X Plus 1.5B flow cell using paired-end sequencing. The samples yielded 277 to 354 million pass-filter reads, with more than 91% of bases above a quality score of Q30. Reads were trimmed for adapters and low-quality bases using Cutadapt version 4.4 ^50^ and aligned to the human reference genome (hg38) and Gencode 30 annotation using STAR in single-pass mode ^51^. The average mapping rate across all samples was 95%, with unique alignment above 86% and 3.09-5.05% unmapped reads. Mapping statistics were calculated using Picard software. The samples contained 1.47% ribosomal bases; coding bases ranged from 64-67%, UTR bases from 28-30%, and mRNA bases from 93-96% across all samples. Library complexity was measured in terms of unique fragments in the mapped reads using Picard’s MarkDuplicates utility, with samples containing 61-66% non-duplicate reads. Gene expression quantification was performed for all samples using RSEM ^52^. Differential gene expression analysis was performed using limma-voom ^53^ as implemented in eVITTA, a web-based visualization and inference toolbox for transcriptome analysis ^54^. Alternative splicing events were quantified using rMATS (v4.3.0) ^30^, which detects and statistically evaluates five categories of alternative splicing events - skipped exons (SE), alternative 5′ splice sites (A5SS), alternative 3′ splice sites (A3SS), mutually exclusive exons (MXE), and retained introns (RI) - from replicate RNA-seq data. Differential splicing events from all five categories were filtered by FDR < 0.05 and |ΔPSI| ≥ 0.10, collapsed to the gene level, and ranked by ascending FDR and descending |ΔPSI| to select the top 10 genes for downstream analysis. QIAGEN Ingenuity Pathway Analysis (IPA) ^25^ was performed by importing the differential gene expression list comparing Mock to SeV-infected WT cells. Analysis-ready molecules were defined using an expression log ratio cutoff of |logFC| ≥ 2.0 (with ≤2.0 as Down and ≥2.0 as Up), yielding 452 analysis-ready molecules (11 Down and 441 Up). Core analyses were performed using the Ingenuity Knowledge Base (Genes Only) as the reference set. Enrichment significance was determined by Fisher’s exact test, and activation states of upstream regulators and pathways were inferred using IPA’s z-score algorithm. A graphical summary was generated directly from the expression dataset to provide an overview of the most significantly enriched pathways, upstream regulators, and biological functions, without the addition of external nodes. A custom interaction network was constructed in which NONO was manually added to the network, and IPA’s Molecule Activity Predictor (MAP) tool was enabled to simulate the directional consequences of NONO activity on connected molecules, including IFNB1 and IFN family cytokines. Differential expression values from the DEG dataset were overlaid onto the resulting network to visualize predicted directional changes and connectivity among these molecules in the context of antiviral and IFN-mediated signaling.

### ATAC-seq data processing

ATAC-seq sample and library preparation was performed on 100,000 fresh cells using the ATAC-seq Kit (Active Motif) according to the manufacturer’s instructions. ATAC-seq libraries were sequenced on a NovaSeq X Plus 1.5B flow cell using paired-end sequencing. The samples yielded 197 to 421 million pass-filter reads, with more than 92% of bases above a quality score of Q30. ATAC-seq reads were processed using the chrom-seek (2.0.0) pipeline. In brief, reads were trimmed with Cutadapt version 4.4 ^50^ and then aligned the human GRCh38.p12 genome using BWA version 0.7.17 ^55^. All reads aligning to the Encode hg38 v2 blacklist regions ^56^ were identified and removed with Picard SamToFastq. Reads with a mapQ score less than 6 were removed with SAMtools version 1.17 ^57^ and PCR duplicates were removed with Picard MarkDuplicates. Data was converted into bigwigs for viewing and normalized by reads per genomic content (RPGC) using deepTools version 3.5.5 ^58^. Averaged bigwigs as well as TSS-associated heatmaps and profile plots were also created using deepTools and ggplot2 version 4.0.3. Peaks were called using Genrich version 0.6. Consensus peaks across all conditions and associated PCA based upon peak intensities was calculated using DiffBind v2 ^59^. Differential peaks were called using DiffBind v2 and its Deseq2 ^60^ differential caller with default parameters. Peaks were then annotated to nearest TSS using UROPA version 4.0.2 ^61^ and Gencode Release 28. Motif analysis was performed on differential peaks using HOMER v 4.11.1 ^62^ with default parameters. Volcano plot was created using GraphPad Prism 10. Example region plots were created using karyoploteR version 1.36.0.

## QUANTIFICATION AND STATISTICAL ANALYSIS

All data are presented as means ± SD and analyzed using GraphPad PRISM software (version 10.5.0 GraphPad Software). Welch’s t-test or two-way ANOVA with Sidak’s or Tukey’s multiple comparisons were used. *p < 0.05; **p < 0.01; ***p < 0.001; ****p < 0.0001.

### Key resources table

| REAGENT or RESOURCE | SOURCE | IDENTIFIER |
| --- | --- | --- |
| Antibodies |  |  |
| anti-NONO (2 $\mu\text{g}/\text{ChIP}$) | Proteintech | Cat# 11058-1-AP;<br>RRID: AB_2152167 |
| anti-NONO (1:1000) | Invitrogen | Cat# MA3-2024;<br>RRID: AB_2152170 |
| anti-pTBK1 (S172) (1:1000) | Cell Signaling<br>Technology | Cat# 5483; RRID:<br>AB_10693472 |
| anti-TBK1 (1:1000) | Cell Signaling<br>Technology | Cat# 51872; RRID:<br>AB_2799403 |
| anti-pIKKε (S172) (1:1000) | Cell Signaling<br>Technology | Cat# 8766; RRID:<br>AB_2737061 |
| anti-IKKε (1:1000) | Cell Signaling<br>Technology | Cat# 2690; RRID:<br>AB_915926 |
| anti-pIRF3 (S396) (1:1000) | Cell Signaling<br>Technology | Cat# 4947; RRID<br>:AB_823547 |
| anti-IRF3 (1:1000) | Cell Signaling<br>Technology | Cat# 11904; RRID:<br>AB_2722521 |
| anti-IRF3 (1:1000) | Cell Signaling<br>Technology | Cat# 10949; RRID:<br>AB_2797733 |
| anti-pSTAT1 (Y701) (1:1000) | Cell Signaling<br>Technology | Cat# 9167; RRID:<br>AB_561284 |
| anti-STAT1 (1:1000) | Cell Signaling<br>Technology | Cat# 14994; RRID:<br>AB_2737027 |
| anti-pSTAT2 (Y690) (1:1000) | Cell Signaling<br>Technology | Cat# 88410; RRID:<br>AB_2800123 |
| anti-STAT2 (1:1000) | Cell Signaling<br>Technology | Cat# 72604; RRID:<br>AB_2799824 |
| anti-pNFκB p65 (S536) (1:1000) | Cell Signaling<br>Technology | Cat# 3033; RRID:<br>AB_331284 |
| anti-NFκB p65 (1:1000) | Cell Signaling<br>Technology | Cat# 6956; RRID:<br>AB_10828935 |
| anti-IRF1 (1:1000) | Cell Signaling<br>Technology | Cat# 8478;<br>RRID:AB_10949108 |
| anti-FRA1 (1:1000) | Cell Signaling<br>Technology | Cat# 5281;<br>RRID:AB_10557418 |
| anti-ATF2 (1:1000) | Cell Signaling<br>Technology | Cat# 35031;<br>RRID:AB_2799069 |
| anti-SFPQ (1:5000) | Proteintech | Cat# 67129-1-Ig;<br>RRID:AB_2882428 |
| anti-PSPC1 (1:500) | Santa Cruz<br>Biotechnology | Cat# sc374181;<br>RRID:AB_10989076 |
| anti-West Nile virus NS5 (1:500) | GeneTex | Cat# GTX131961;<br>RRID: AB_2800433 |
| anti-Zika virus NS5 (1:1000) | GeneTex | Cat# GTX133312;<br>RRID: AB_2750559 |
| anti-Yellow Fever virus NS3 (1:1000) | GeneTex | Cat# GTX133959;<br>RRID: AB_2887155 |
| anti-β-Actin (1:5000) | Sigma-Aldrich | Cat# A5441; RRID:<br>AB_476744 |
| anti-β-Tubulin (1:2000) | Sigma-Aldrich | Cat# T8328; RRID:<br>AB_1844090 |
| anti-Lamin A/C (1:1000) | Santa Cruz<br>Biotechnology | Cat# sc-376248;<br>RRID: AB_10991536 |
| anti-RNA Pol II (2 μg/ChIP) | Active Motif | Cat# 91151; RRID:<br>AB_2793789 |
| anti-RNA Pol II CTD (1:1000) | Cell Signaling<br>Technology | Cat# 2629;<br>RRID:AB_2167468 |
| anti-Histone H3 (1:1000) | Cell Signaling<br>Technology | Cat# 14269;<br>RRID:AB_2756816 |
| anti-Tri-Methyl-Histone H3 (Lys4) (2 μg/ChIP) | Cell Signaling<br>Technology | Cat# 9751; RRID:<br>AB_2616028 |
| anti-Tri-Methyl-Histone H3 (Lys27) (2 μg/ChIP) | Cell Signaling<br>Technology | Cat# 9733; RRID:<br>AB_2616029 |
| anti-Rabbit IgG Isotype Control (2 µg/ChIP) | Cell Signaling Technology | Cat# 2729; RRID: AB_1031062 |
| anti-Mouse IgG1 Isotype Control (2 µg/ChIP) | Cell Signaling Technology | Cat# 5415; RRID: AB_10829607 |
| anti-Rabbit IgG (whole molecule)-peroxidase antibody produced in goat (1:5000) | Sigma-Aldrich | Cat# A6154; RRID: AB_258284 |
| anti-Mouse IgG (whole molecule)-peroxidase antibody produced in goat (1:4000) | Sigma-Aldrich | Cat# A4416; RRID: AB_258167 |
| Bacterial and virus strains |  |  |
| Zika virus (Dakar) | World Reference Center of Emerging Viruses and Arboviruses (WRCEVA) | N/A |
| Yellow fever virus (Asibi) | World Reference Center of Emerging Viruses and Arboviruses (WRCEVA) | N/A |
| Tick-borne encephalitis virus (Sofjin) | World Reference Center of Emerging Viruses and Arboviruses (WRCEVA) | N/A |
| Kyasanur Forest disease virus (P9605) | World Reference Center of Emerging Viruses and Arboviruses (WRCEVA) | N/A |
| Omsk hemorrhagic fever virus (Guriev) | World Reference Center of Emerging Viruses and Arboviruses (WRCEVA) | N/A |
| West Nile virus (Bird 114) | BEI Resources | Cat# NR-9538 |
| West Nile virus (NY99) | BEI Resources | Cat# NR-158 |
| Yellow fever virus (17D) | BEI Resources | Cat# NR-116 |
| Dengue virus serotype 2 (NGC) | BEI Resources | Cat# NR-84 |
| Sendai virus (Cantell) | AVS Bio | Cat# 10100774 |
| Chemicals, peptides, and recombinant proteins |  |  |
| Lipofectamine 2000 | Invitrogen | Cat# 11668019 |
| Lipofectamine RNAiMAX | Invitrogen | Cat# 13778150 |
| Poly(I:C) LMW | InvivoGen | Cat# tlrl-picw |
| Poly(I:C) HMW | InvivoGen | Cat# tlrl-pic |
| 3p-hpRNA | InvivoGen | Cat# tlrl-hprna |
| Poly(dA:dT) naked | InvivoGen | Cat# tlrl-patn |
| Human IFN-Beta 1a, mammalian, carrier-free | PBL Assay Science | Cat# 11410 |
| Ruxolitinib | InvivoGen | Cat# tlrl-rux-3 |
| Dimethyl sulfoxide | Sigma-Aldrich | Cat# D2653 |
| N-ethylmaleimide | Sigma-Aldrich | Cat# E3876 |
| Protein A/G agarose beads | Thermo Scientific | Cat# 20421 |
| cOmplete, Mini, EDTA-free Protease Inhibitor Cocktail | Roche | Cat# 11836170001 |
| ECL Western Blotting Substrate | Thermo Scientific | Cat# 32106 |
| SuperSignal West Femto Maximum Sensitivity Substrate | Thermo Scientific | Cat# 34096 |
| Critical commercial assays |  |  |
| Trans-Blot Turbo Mini 0.2 µm PVDF Transfer Packs | Bio-Rad | Cat# 1704156 |
| RNeasy Plus Mini Kit | Qiagen | Cat# 74136 |
| SuperScript VILO Master Mix | Invitrogen | Cat# 11755250 |
| SsoAdvanced Universal SYBR Green Supermix | Bio-Rad | Cat# 1725275 |
| SimpleChIP Plus Sonication Chromatin IP Kit | Cell Signaling Technology | Cat# 56383 |
| SimpleChIP Universal qPCR Master Mix | Cell Signaling Technology | Cat# 88989 |
| ATAC-seq Kit | Active Motif | Cat# 53150 |
| NEBNext Ultra II DNA Library Prep Kit | New England Biolabs | Cat# E7645S |
| LumiKine Xpress hIFN-β 2.0 ELISA Kit | Invivogen | Cat# luex-hifnbv3 |
| SF Cell Line 4D-Nucleofector X Kit S | Lonza | Cat# V4XC-2032 |
| NE-PER Nuclear and Cytoplasmic Extraction Kit | Thermo Scientific | Cat# 78833 |
| Subcellular Protein Fractionation Kit for Cultured Cells | Thermo Scientific | Cat# 78840 |
| Deposited data |  |  |
| RNA-seq data | This paper | GEO: GSE335258 |
| ATAC-seq data | This paper | GEO: GSE335260 |
| Experimental models: Cell lines |  |  |
| A549 (WT and *NONO* KO) | ATCC and Sonja M. Best | CCL-185 and this paper |
| Vero | ATCC | CCL-81 |
| Oligonucleotides |  |  |
| Primers for qRT-PCR used in this study | This paper and OriGene | Table S1 |
| Primers for ChIP-PCR used in this study | This paper and Cell Signaling Technology | Table S1 |
| ON-TARGETplus Non-targeting Control Pool | Horizon (Dharmacon) | Cat# D-001810-10-05 |
| ON-TARGETplus Human *NONO* siRNA | Horizon (Dharmacon) | Cat# L-007756-01-0005 |
| Software and algorithms |  |  |
| GraphPad Prism 10 | GraphPad Software, LLC | https://www.graphpad.com/ |
| rMATS (4.3.0) | SourceForge | https://rnaseq-mats.sourceforge.io/ |
| QIAGEN Ingenuity Pathway Analysis (IPA) content version 153384343 | QIAGEN | https://www.qiagen.com/ |
| chrom-seek (2.0.0) | GitHub, Inc | https://github.com/OpenOmics/chrom-seek/releases |
| Picard SamToFastq | GitHub, Inc | https://broadinstitute.github.io/picard/ |
| Genrich (0.6) | GitHub, Inc | https://github.com/jsh58/Genrich |
| deepTools (3.5.5) | GitHub, Inc | https://github.com/deeptools/deeptools/releases |
| ggplot2 (4.0.3) | GitHub, Inc | https://github.com/tidyverse/ggplot2 |
| HOMER (4.11.1) | Benner Lab (UCSD) | http://homer.ucsd.edu/homer/ |
| karyoploteR (1.36.0) | GitHub, Inc | https://github.com/benratgel/karyoploteR |
| Other |  |  |
| GloMax Explorer Multimode Microplate Reader | Promega | Cat# GM3500 |
| iBright FL1500 Imaging System | Invitrogen | Cat# A44241 |
| Trans-Blot Turbo transfer system | Bio-Rad | Cat# 1704150 |
| QuantStudio 7 Pro Real-Time PCR System, 384-well | Applied Biosystems | Cat# A43185 |
| Q800R3 Sonicator | Qsonica | Cat# Q800R3-110 |
| 4D-Nucleofector X Unit | Lonza | Cat# AAF-1003X |

**Table S1:**
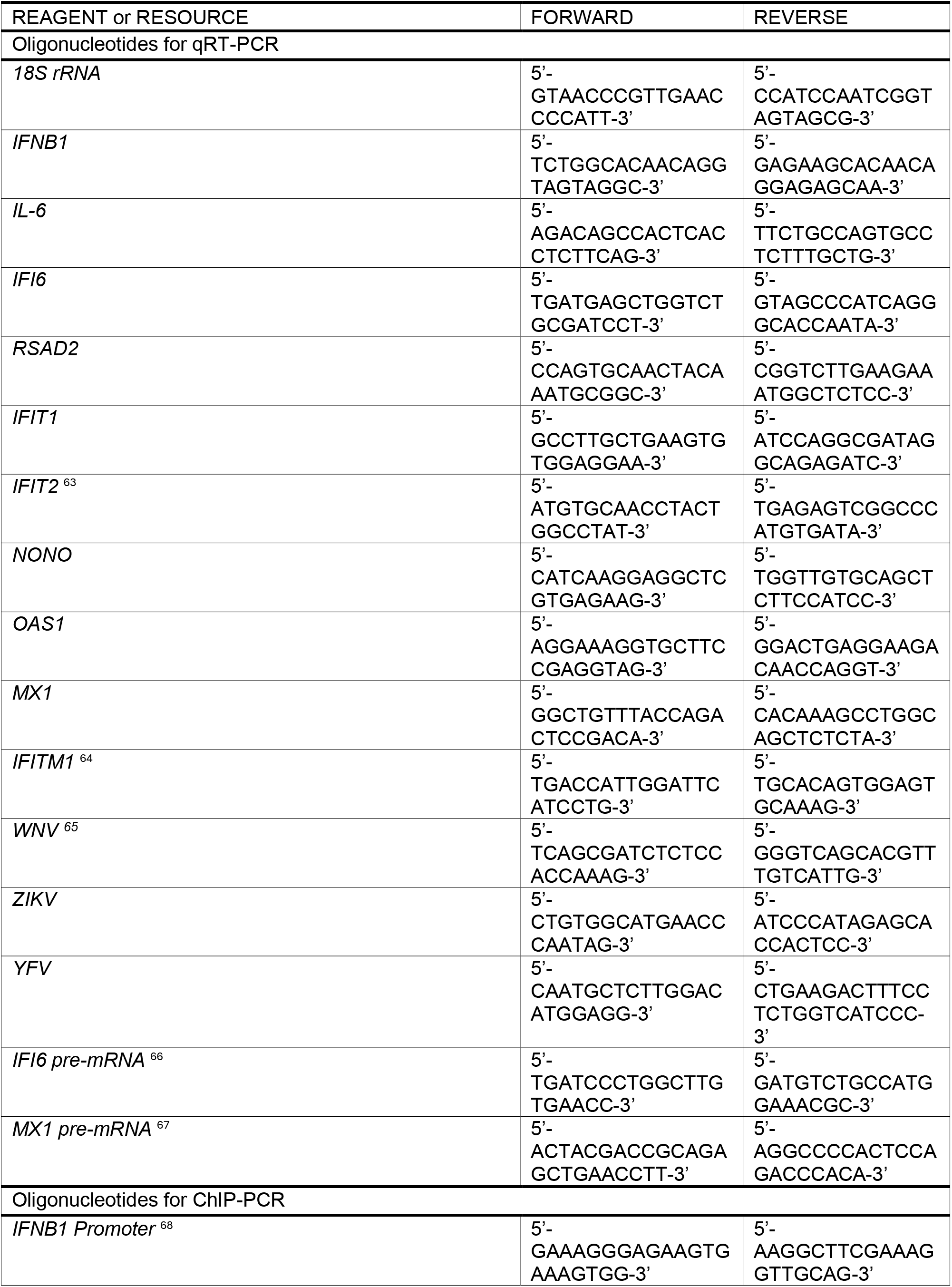
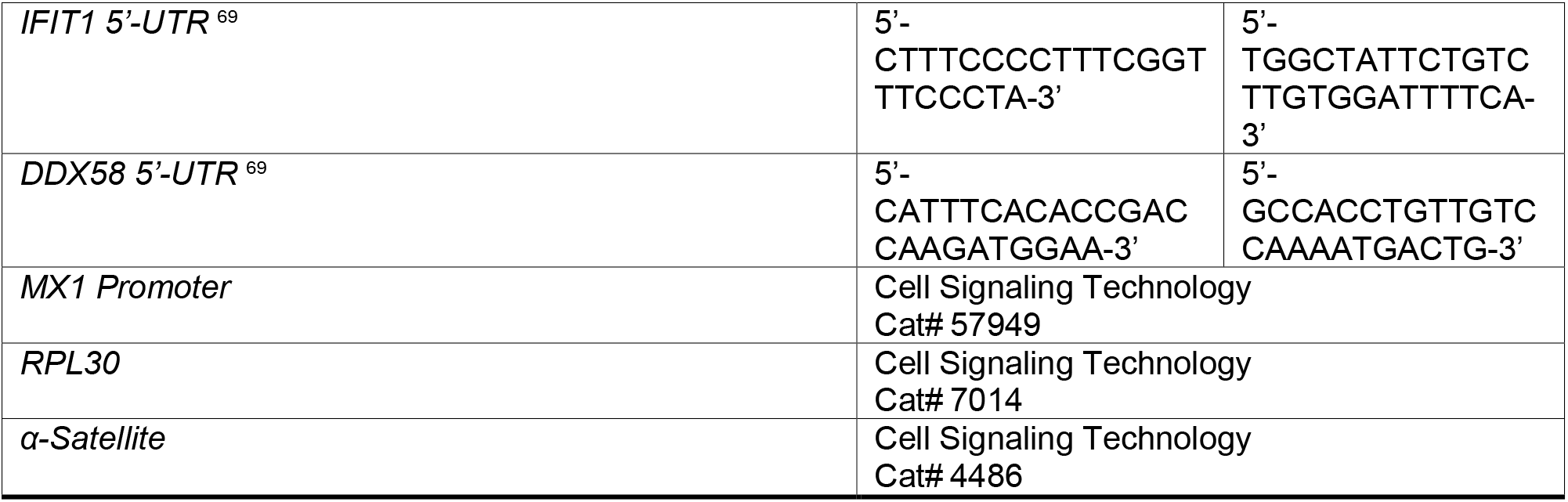
Primer sequences.

---

**Figure S1.**
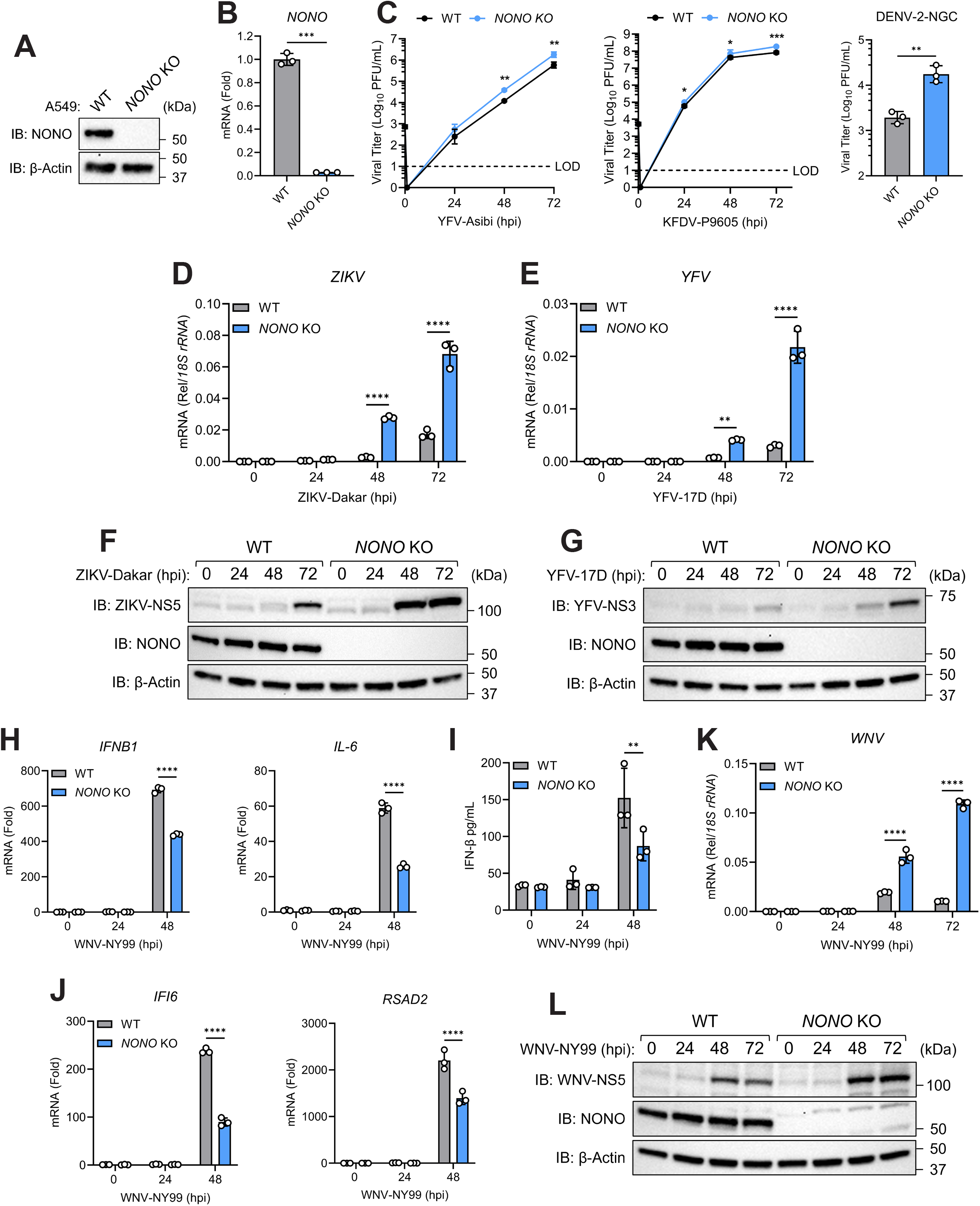
NONO loss enhances orthoflavivirus replication. (A-B) CRISPR knockout efficiency of (A) NONO protein (immunoblot) and (B) *NONO* mRNA (qRT-PCR). (C) Infectious viral titers from WT or *NONO* KO A549 cells infected with yellow fever virus (YFV) Asibi MOI=0.001, Kyasanur Forest disease virus (KFDV) MOI=0.01, and Dengue virus serotype 2 (DENV-2) NGC MOI=1 at 48 hpi (plaque assay). (D-E) *ZIKV* and *YFV-17D* expression from infected WT or *NONO* KO A549 cells from (A) (qRT-PCR). (F-G) Levels of ZIKV-NS5 and YFV-17D-NS3 protein from infected WT or *NONO* KO A549 cells from (A) (immunoblot). (H-L) WT or *NONO* KO A549 cells infected with WNV-NY99 MOI=0.0001. (H) *IFN-β* and *IL-6* expression (qRT-PCR). (I) IFN-β protein in supernatants (ELISA). (J) *IFI6* and *RSAD2* expression (qRT-PCR). (K) *WNV* expression (qRT-PCR). (L) Levels of WNV-NS5 protein (immunoblot). Data are expressed as means (n=3) ± SD, *p < 0.05, **p < 0.01, ***p < 0.001, ****p < 0.0001 (Student’s t-test or two-way ANOVA with Sidak’s multiple comparisons) and are representative of 2-3 independent experiments.

**Figure S2.**
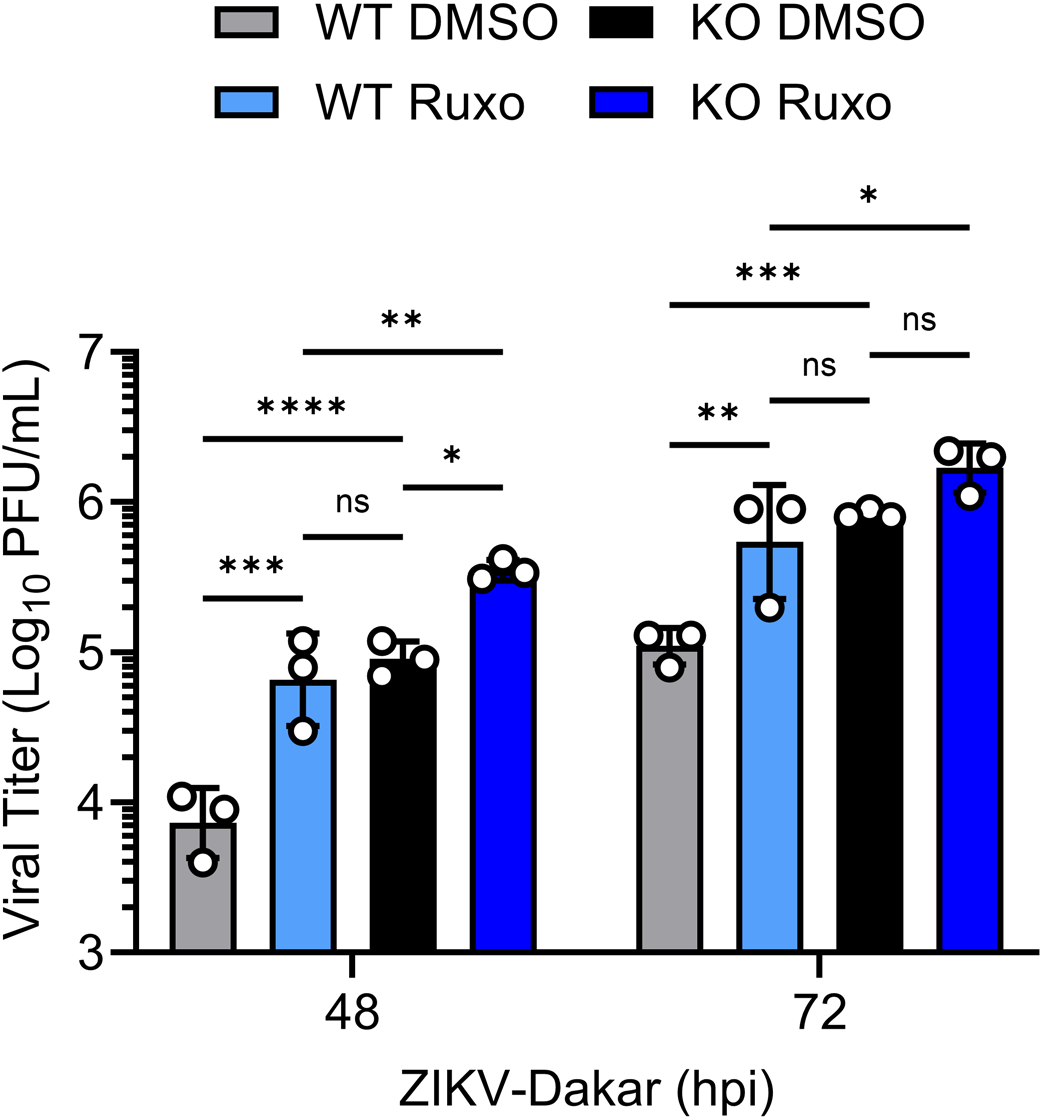
NONO is critically required for establishing an effective antiviral response. Infectious viral titers from WT or *NONO* KO A549 cells pretreated with Ruxolitinib (10 μg/mL) or DMSO carrier for 1 hour before infection with Zika virus (ZIKV) Dakar MOI=0.01 (plaque assay). Data are expressed as means (n=3) ± SD, *p < 0.05, **p < 0.01, ***p < 0.001, ****p < 0.0001 (two-way ANOVA with Tukey’s multiple comparisons) and are representative of 2-3 independent experiments.

**Figure S3.**
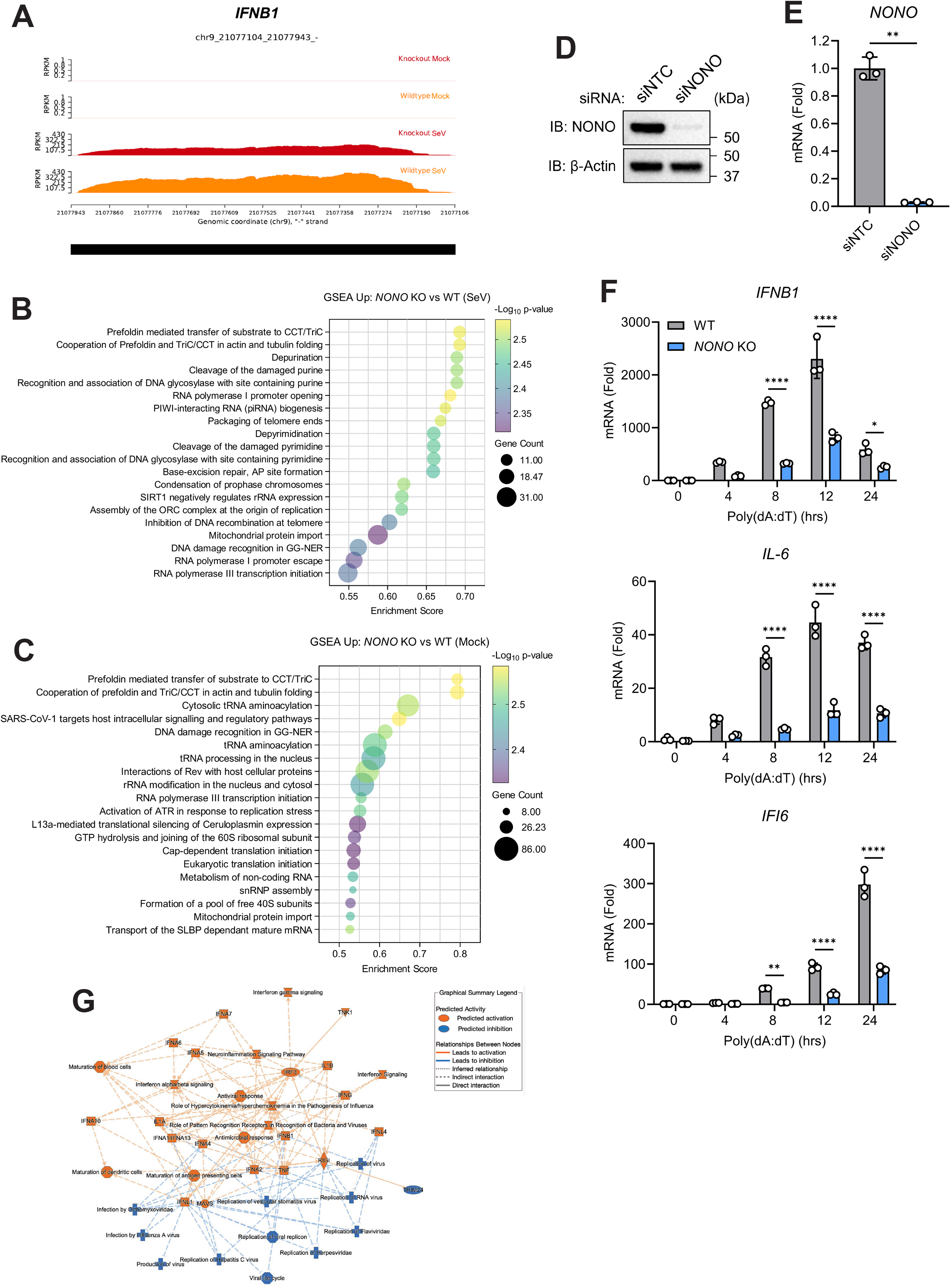
NONO broadly promotes innate immune responses. (A-C) WT or *NONO* KO A549 cells infected with SeV (100 HAU/mL) for 8 hours. (A) Reads Per Kilobase Million (RPKM) for *IFN-β*. (B-C) GSEA terms and pathways of upregulated transcripts from SeV-infected (B) and mock (C) samples. (D-E) siRNA silencing efficiency of (D) NONO protein (immunoblot) and (E) *NONO* mRNA (qRT- PCR). (F) *IFN-β*, *IL-6*, and *IFI6* expression from WT or *NONO* KO A549 cells stimulated with Poly(dA:dT) (1 μg/mL) (qRT-PCR). (G) A graphical summary using Ingenuity Pathway Analysis (IPA) performed by importing the differential gene expression list comparing Mock to SeV-infected WT cells visualizes the most significantly enriched pathways, upstream regulators, and biological functions, without the addition of external nodes. Data are expressed as means (n=3) ± SD, *p < 0.05, **p < 0.01, ****p < 0.0001 (Student’s t-test or two-way ANOVA with Sidak’s multiple comparisons) and are representative of 2-3 independent experiments.

**Figure S4.**
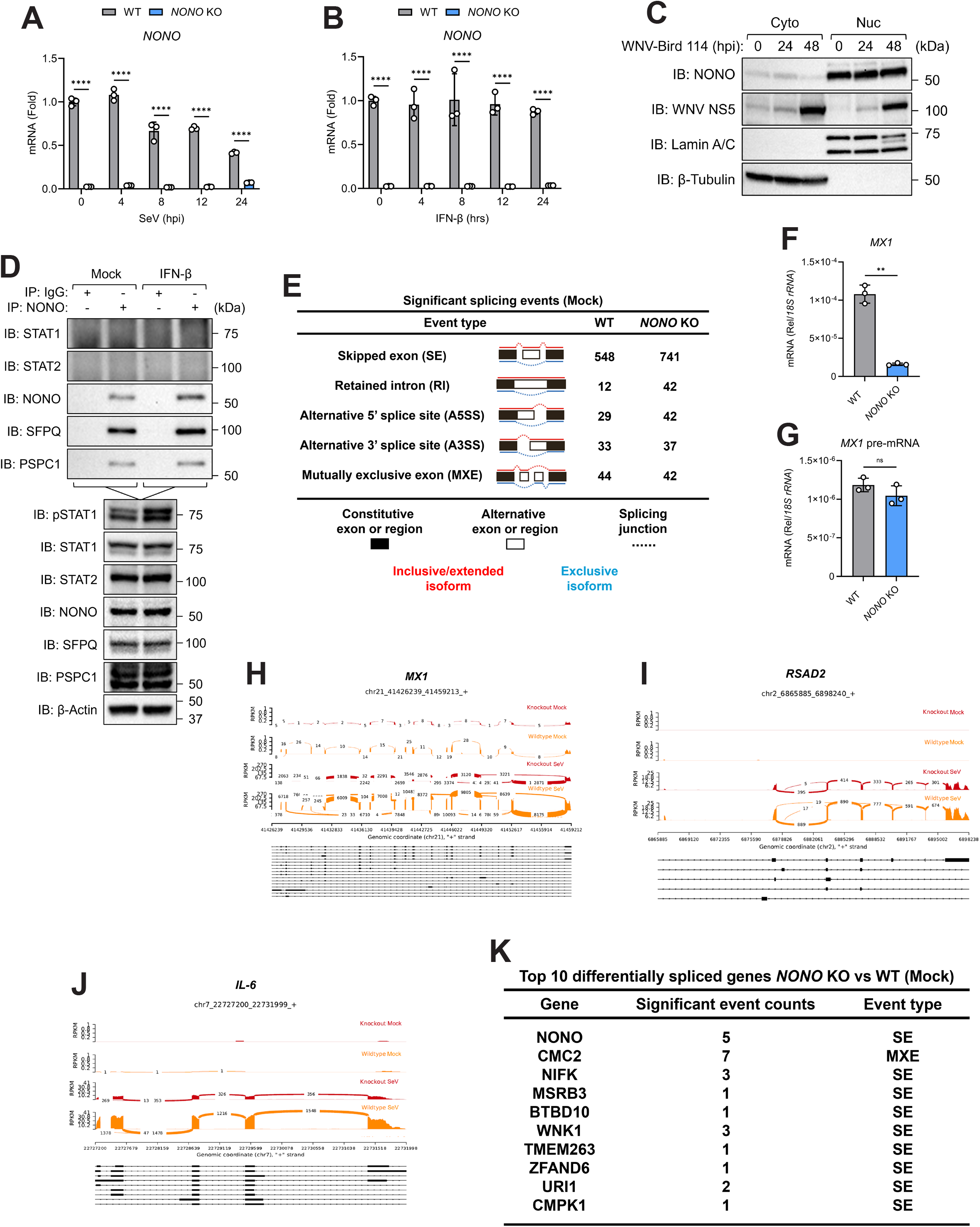
NONO augments innate immunity specifically at the transcriptional level. (A) *NONO* expression from WT or *NONO* KO A549 cells infected with SeV (100 HAU/mL). (B) *NONO* expression from WT or *NONO* KO A549 cells stimulated with IFN-β (100 IU/mL). (C) Subcellular fractionation of NONO and WNV-NS5 protein from WT A549 cells infected with WNV-Bird 114 MOI=0.0001 (immunoblot). (D) Endogenous co-immunoprecipitation of NONO from WT A549 cells stimulated with IFN-β (1000 IU/mL) for 8 hours. (E) rMATS alternative splicing events between WT and *NONO* KO A549 cells. (F-G) Mature and pre- mRNA expression for *MX1* from WT or *NONO* KO A549 cells stimulated with IFN-β (100 IU/mL) for 8 hours (qRT-PCR). (H-J) Sashimi plots for *MX1*, *RSAD2*, and *IL-6* from WT or *NONO* KO A549 cells mock or SeV infected (100 HAU/mL) for 8 hours. (K) rMATS top 10 differentially spliced genes from WT and *NONO* KO A549 cells. Data are expressed as means (n=3) ± SD, **p < 0.01, ****p < 0.0001 (two-way ANOVA with Sidak’s multiple comparisons or Student’s t-test) and are representative of 2-3 independent experiments.

**Figure S5.**
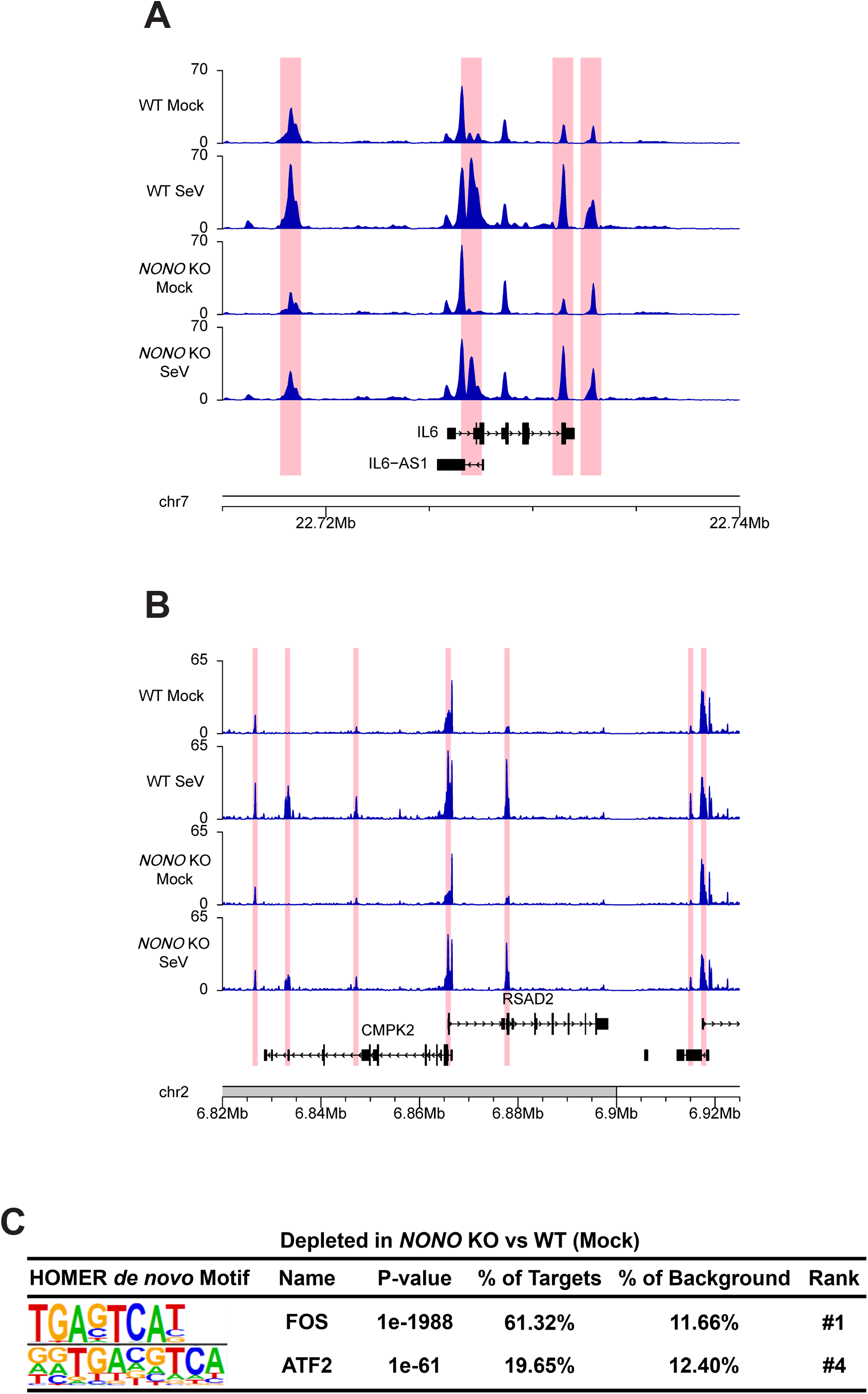
NONO enhances chromatin accessibility of innate immune genes. (A-C) ATAC-seq of WT or *NONO* KO A549 cells infected with SeV (100 HAU/mL) for 8 hours (n=4 per group). (A-B) Genome browser view of open chromatin peaks for *IL-6*, *IL6-AS1*, *RSAD2*, and *CMPK2* with highlighted portions indicating regions with significantly less available chromatin in the SeV-infected *NONO* KO compared to the SeV-infected WT. (C) HOMER *de novo* motif analysis for depleted motifs in mock *NONO* KO compared to WT samples.

**Figure S6.**
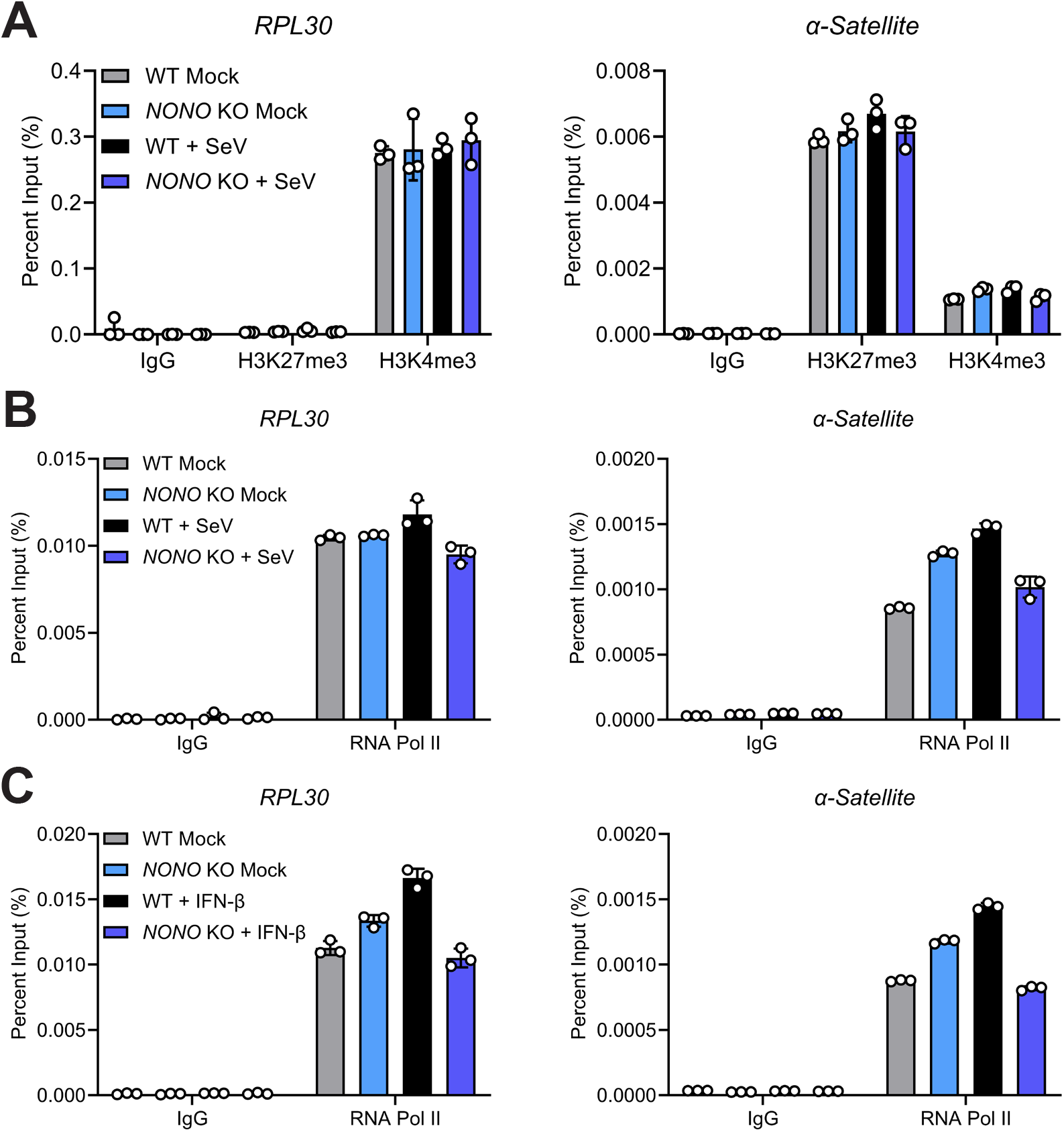
NONO loss impairs RNA Pol II binding to innate immune gene promoters. (A-B) WT or *NONO* KO A549 cells infected with SeV (100 HAU/mL) for 8 hours. (A) ChIP-qPCR showing H3K27me3 and H3K4me3 histone modifications at *RPL30* and *α-Satellite*. (B) ChIP-qPCR showing RNA-Pol II occupancy at *RPL30* and *α-Satellite*. (C) ChIP-qPCR showing RNA-Pol II occupancy at *RPL30* and *α-Satellite* for WT or *NONO* KO A549 cells stimulated with IFN-β (1000 IU/mL) for 8 hours. Data are expressed as means (n=3) ± SD and are representative of 2-3 independent experiments.

## Notes

### Competing Interest Statement

The authors have declared no competing interest.

